# ATF-4 and hydrogen sulfide signalling mediate longevity from inhibition of translation or mTORC1

**DOI:** 10.1101/2020.11.02.364703

**Authors:** Cyril Statzer, Jin Meng, Richard Venz, Monet Bland, Stacey Robida-Stubbs, Krina Patel, Dunja Petrovic, Raffaella Emsley, Pengpeng Liu, Ianessa Morantte, Cole Haynes, William B. Mair, Alban Longchamp, Milos Filipovic, T. Keith Blackwell, Collin Y. Ewald

**Affiliations:** Eidgenössische Technische Hochschule Zürich, Department of Health Sciences and Technology, Institute of Translational Medicine, Schwerzenbach, Switzerland; Department of Genetics, Harvard Medical School, Boston, MA, United States; Joslin Diabetes Center, Research Division, Boston, MA, United States; Department of Vascular Surgery, Centre Hospitalier Universitaire Vaudois and University of Lausanne, Lausanne, Switzerland; Leibniz-Institut für Analytische Wissenschaften-ISAS-e.V., Dortmund, Germany; Department of Molecular, Cell and Cancer Biology, University of Massachusetts Medical School, Worcester, MA, U.S.A; Department of Genetics and Complex Diseases, Harvard School of Public Health, 665 Huntington Avenue, Boston, MA, U.S.A

**Keywords:** mRNA translation, cystathionine gamma-lyase, H_2_S, ageing, mTORC1, integrated stress response, *C*. *elegans*

## Abstract

Inhibition of the master growth regulator mTORC1 (mechanistic target of rapamycin complex 1) slows ageing across phyla, in part by reducing protein synthesis. Various stresses globally suppress protein synthesis through the integrated stress response (ISR), resulting in preferential translation of the transcription factor ATF-4. Here we show in *C. elegans* that inhibition of translation or mTORC1 increases ATF-4 expression, and that ATF-4 mediates longevity under these conditions independently of ISR signalling. ATF-4 promotes longevity by activating canonical anti-ageing mechanisms, but also by elevating expression of the transsulfuration enzyme CTH-2 to increase hydrogen sulfide (H_2_S) production. This H_2_S boost increases protein persulfidation, a protective modification of redox-reactive cysteines. The ATF-4/CTH-2/H_2_S pathway also mediates longevity and increased stress resistance from mTORC1 suppression. Increasing H_2_S levels, or enhancing mechanisms that H_2_S influences through persulfidation, may represent promising strategies for mobilising therapeutic benefits of the ISR, translation suppression, or mTORC1 inhibition.

## Introduction

Over the last three decades, genetic and phenotypic analyses of ageing have revealed that across eukaryotes, lifespan can be extended by inhibition of mechanisms that promote growth and proliferation^1–3^. Prominent among these is the serine/threonine kinase complex mTORC1, which coordinates the activity of multiple growth-related processes in response to growth factor and nutrient signals^2–4^. mTORC1 activity can be reduced by genetic perturbations, dietary restriction (DR), or pharmacological interventions such as rapamycin, an mTORC1 inhibitor that increases lifespan from yeast to mice^2, 3^. However, while rapamycin represents a very exciting paradigm for anti-ageing pharmacology, mTORC1 suppression has wide-ranging effects on the organism^2, 3^. Rapamycin is used clinically as an immunosuppressant, and mTORC1 broadly affects metabolism and supports the synthesis of proteins, nucleic acids, and lipids^2–4^. Elucidation of specific mechanisms through which mTORC1 influences longevity is critical not only for understanding the biology of ageing and longevity, but also for the development of molecularly targeted anti-ageing therapies that maintain health.

mTORC1 increases the rates at which numerous different mRNAs are translated, and a hallmark of mTORC1 inhibition is a reduction in overall protein synthesis^2, 3^. Work in the model organisms *C. elegans* and *Drosophila* indicates that lifespan extension from mTORC1 inhibition is mediated in part through this global decrease in translation^2, 5, 6^. Furthermore, in *C. elegans* suppression of translation is sufficient to increase both lifespan and stress resistance^7–12^. A mechanistic understanding of how mRNA translation levels affect longevity should therefore provide mechanistic insights into how mTORC1 influences lifespan.

Suppression of new protein synthesis is also an important mechanism through which cells protect themselves from stressful conditions that include nutrient deprivation, and thermal-, oxidative-, and endoplasmic reticulum (ER) stress^13, 14^. Those stresses induce the evolutionarily conserved ISR by activating kinases that phosphorylate and inhibit the translation initiation factor subunit eIF2α, thereby imposing a broad reduction in cap-dependent mRNA translation^13, 14^. This suppression of translation leads in turn to preferential translation of the activating transcription factor ATF4, which mobilizes stress defense mechanisms to reestablish homeostasis^13, 14^. Although the ISR has important protective functions, its effects on longevity and health are complex. In *C. elegans*, genetic or pharmacological interventions that impair the ISR extend lifespan by inducing preferential translation of selective mRNAs^15^. In older mice, pharmacological ISR inhibition enhances memory and cognition by allowing protein synthesis to be maintained^14, 16^. On the other hand, in *S. cerevisiae* the ATF4 ortholog Gcn4 promotes longevity^17, 18^, and in *C. elegans* hexosamine pathway activation enhances proteostasis through the ISR and ATF-4^19^. The effects of ATF4 on metazoan longevity have not been explored.

Here we have investigated whether and how ATF-4 affects lifespan in *C. elegans*. We find that ATF-4 but not upstream ISR signalling is required for longevity induced by conditions that inhibit protein synthesis. Importantly, ATF-4 is a pro-longevity factor that extends lifespan when overexpressed on its own. ATF-4 increases lifespan not only by enhancing canonical anti-ageing mechanisms, but also by inducing transsulfuration enzyme-mediated H_2_S production and therefore levels of protein persulfidation. Importantly, the anti-ageing benefits of mTORC1 suppression depend upon this ATF-4-induced increase in H_2_S production, further supporting the idea that they derive from lower translation rates and suggesting that increases in ATF-4 and H_2_S levels may recapitulate these benefits.

## Results

### ATF-4 responds to translation suppression to increase *C. elegans* lifespan

We investigated whether *C. elegans atf-4* is regulated similarly to mammalian ATF4 at the level of mRNA translation. In mammals, 2-3 small upstream open reading frames (uORFs) within the ATF4 5’ untranslated region (UTR) occupy the translation machinery under normal conditions, inhibiting translation of the downstream ATF4 coding region^14, 20^. By contrast, when translation initiation is impaired by eIF2α phosphorylation, the last uORF is bypassed, and ATF4 is translated preferentially^14, 20^. The *C. elegans atf-4* ortholog (previously named *atf-5*) contains two 5’ UTR uORFs (Fig. 1a; Extended Data Fig. 1a, 1b), deletion of which increases translation of a transgenic reporter^21^, predicting that translation of the *atf-4* mRNA may be increased under conditions of global translation suppression.

**Fig. 1.**
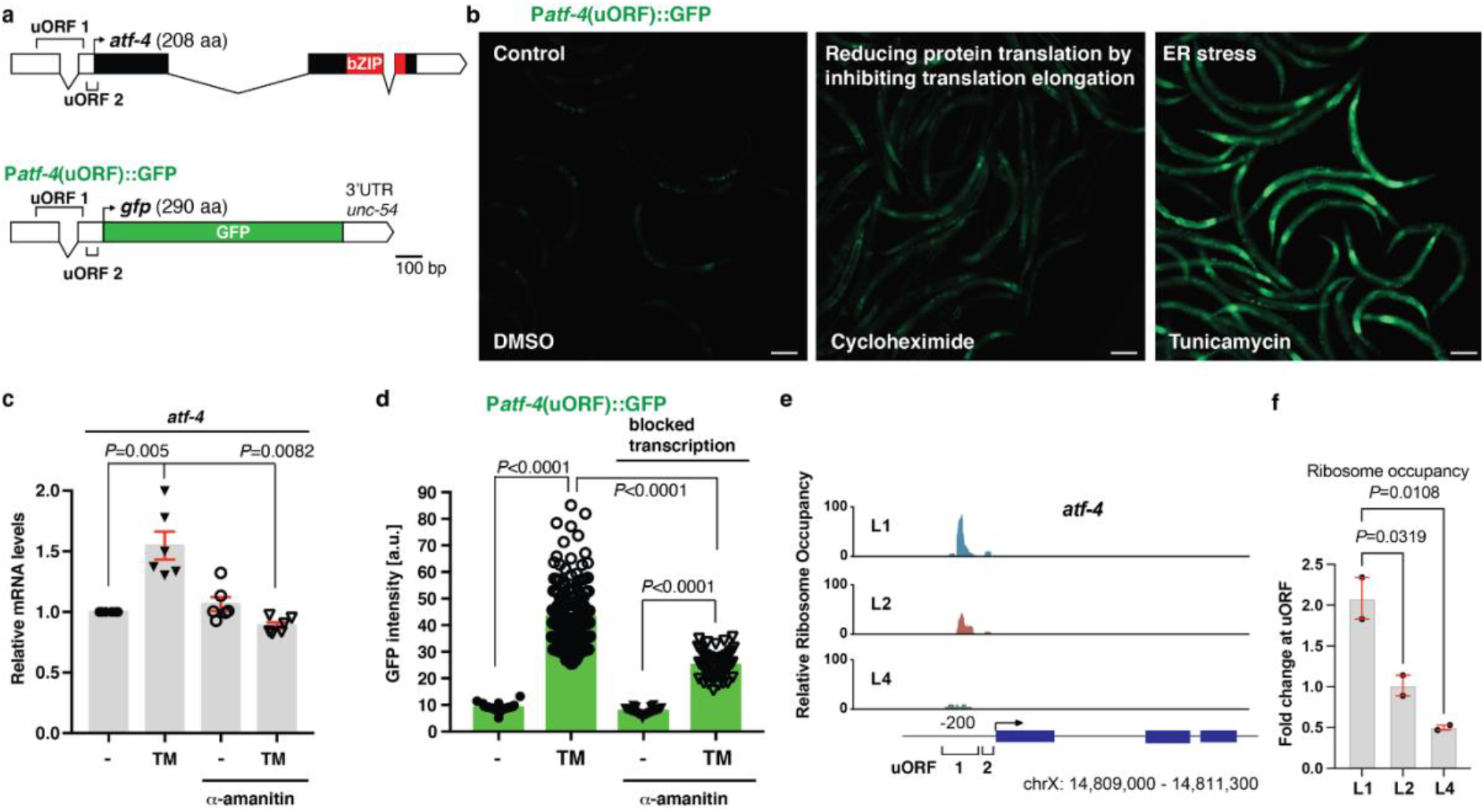
ATF-4 is preferentially translated under conditions of reduced global protein synthesis. **a** Schematic diagram of the *atf-4* mRNA and the P*atf-4*(uORF)::GFP reporter. UTRs are represented as empty boxes, exons as filled boxes, and the basic leucine zipper domain (bZIP) in red. **b** Representative images showing that reducing translation by administering 7.2 mM cycloheximide for 1 hour or 35 μg/ml tunicamycin (TM) for 4 hours increased expression of transgenic P*atf-4*(uORF)::GFP in L4 stage animals. Quantification of GFP fluorescence intensity is shown in **Extended Data Fig. 1c**. Scale bar = 100 μm. **c** A 1 hour pre-treatment with 0.7 μg/ml α-amanitin (RNA Pol II inhibitor) prevented 4 hours of 35 μg/ml TM treatment from increasing *atf-4* mRNA levels in L4 stage animals. Mean ± SEM. Three independent trials, measured in duplicates. *P* values are relative to WT (N2) determined by one sample *t*-test, two-tailed, hypothetical mean of 1. **d** A 1 hour pre-treatment with 0.7 μg/ml α-amanitin did not prevent TM treatment from increasing levels of transgenic P*atf-4*(uORF)::GFP expression in L4 stage animals. Mean ± SEM. n>30 animals, 2 independent trials, One-way ANOVA with post hoc Tukey. **e** Stage-specific ribosome occupancy profiles of the endogenous *atf-4* mRNA, along with quantification of relative uORF occupancy (**f**). Analysis of ribosomal profiling data^55^ revealed a decrease in ribosome occupancy on the endogenous *atf-4* uORFs under unstressed conditions during late larval development. Occupancy profiles were generated by assigning counts to the *atf-4* transcript based on the number of raw reads at each position. Blue boxes indicate the *atf-4* exons. One-way ANOVA post hoc Dunnett’s test.

**Extended Data Fig. 1.**
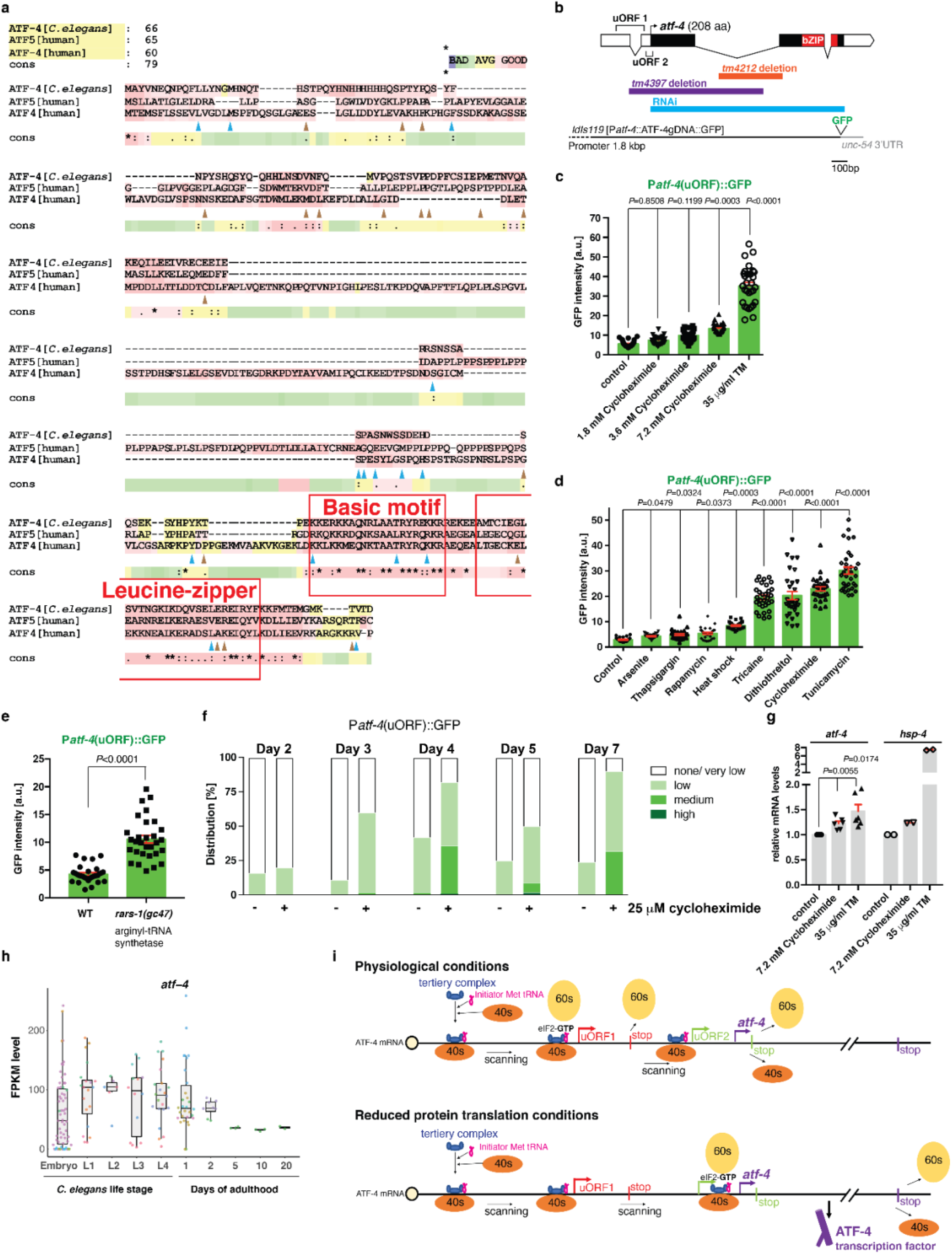
Translational regulation of *C. elegans atf-4* expression. **A** *elegans* ATF-4 (T04C10.4) shares identical amino acids with human ATF4 (blue arrowheads) and ATF5 (brown arrowheads), and its basic DNA binding region is more similar to that of human ATF4. Stars indicate identical amino acids among *C. elegans* ATF-4, human ATF4, and ATF5 (25 in total). Single dots indicate that size or hydropathy is conserved, while double dots indicate that both size and hydropathy are conserved between the corresponding residues. The *C. elegans* ortholog of human ATF5 is ATFS1^56^. **b** Diagram of *atf-4* mRNA, mutations and RNAi clone, and the *Patf-4(uORF)::GFP* transgene. The *atf-4* mRNA has an extensive 5’UTR of 250 nucleotides containing two uORFs, of which uORF1 translates into a 39 amino acids (aa) peptide and uORF2 into a 14 aa peptide. The *tm4397* variation is an 806 base pair (bp) deletion that covers part of the uORF1, the uORF2, the translational start site and the first exon, suggesting that *tm4397* is a putative null allele. **c** Quantification of GFP fluorescence in P*atf-4*(uORF)::GFP transgenic animals at the L4 stage treated either with cycloheximide for 1 hour or TM for 4 hours. Mean ± SEM. One-way ANOVA with post hoc Tukey. **d** Quantification of GFP fluorescence showing the effects on ATF-4 expression of various drug treatments or interventions that reduce mRNA translation. L4 stage animals were treated either with 20 mM arsenite (an inducer of oxidative stress) for 30 min, 200 mM thapsigargin (which induces ER stress by inhibiting the ER Ca^2+^ ATPase) for 4 hours, 100 μM rapamycin (an inhibitor of mTORC1) for overnight, heat shock at 35°C for 30 min, 2% tricaine for 1 hour (which induces ER stress), 10 mM dithiothreitol (which induces reductive ER stress) for 4 hours, 10 mM cycloheximide (an inhibitor of translation elongation) for 1 hour, or 35 μg/ml tunicamycin (a glycosylation inhibitor that induces ER stress) for 4 hours. Mean ± SEM. One-way ANOVA with post hoc Tukey. **e** Nonsense mutation in the arginyl-tRNA synthetase *rars-1(gc47)* increased P*atf-4*(uORF)::GFP expression compared to WT at the L4 stage. Mean ± SEM. Unpaired two-tailed Student’s *t*-test, hypothetical mean of **f** Quantification of GFP intensity in transgenic P*atf-4*(uORF)::GFP animals at different ages showing that 25 uM cycloheximide induced ATF-4 induction. n=2 independent trials. L4 animals were transferred onto plates containing either DMSO or cyclohexmide with FUdR. **g** Quantification of *atf-4* mRNA levels after cycloheximide or TM treatment in L4 stage animals. n=3 independent trials, measured in duplicates. In one trial, *hsp-4* mRNA was assessed as a positive control for ER stress. Mean ± SEM. *P* values relative to control determined by one-sample *t*-test, two-tailed, a hypothetical mean of 1. **h** Expression levels of *atf-4* mRNA plotted as Fragments Per Kilobase of transcript per Million mapped reads (FPKM) during development and ageing. The *atf-4* mRNA expression levels of untreated WT *C. elegans* were retrieved using the RNAseq FPKM Gene Search tool (www.wormbase.org). The boxplots represent the overall expression pattern and the color of the individual dots refer to the 32 individual studies used. Hypothetical working model for *C. elegans atf-4* preferential translation, assuming that its regulation is the same as mammalian *ATF4*^14, 20^. The *C. elegans atf-4* gene encodes two uORFs. After translating the first uORF, the small ribosomal subunit will continue scanning along the ATF4 mRNA. Under non-stressed conditions, *i.e.*, when high amounts of the eIF2-GFP bound Met-tRNA ^Met^ are available, the small ribosomal subunit will readily acquire the eIF2 ternary complex, and the large ribosomal subunit will associate to translate the second uORF. In mammalian ATF4, the ribosome disassociates from the *atf-4* mRNA after translating the last uORF. However, under stress or reduced translational conditions, *i.e.,* low amounts of the eIF2-GFP bound Met-tRNA ^Met^ availability, the association of the large to the small ribosomal subunit is delayed, whereby the inhibitory second uORF is skipped and the re-initiation complex starts to translate the ATF-4 coding region. Phosphorylation of eIF2α subunit inhibits the guanine nucleotide exchange factor eIF2B, which lowers the exchange of the eIF2-GDP to eIF2-GTP and thereby lowers global mRNA translation initiation.

We tested this idea in *C. elegans* that express green fluorescent protein (GFP) driven by the *atf-4* upstream region, including the two uORFs (P*atf-4*(uORF)::GFP, Fig. 1a). P*atf-4*(uORF)::GFP expression was extremely low under unstressed conditions, but increased when translation was suppressed by treatment with the translation elongation blocker cycloheximide, or RNA interference (RNAi)-mediated knockdown of the tRNA synthase *rars-1* (Fig. 1b, Extended Data Fig. 1c, 1d, 1e, 1f). Compounds that induce ER stress and elicit the ISR, including tunicamycin (TM) or dithiothreitol (DTT), also strongly induced P*atf-4*(uORF)::GFP expression (Fig. 1b, Extended Data Fig. 1c, 1d). By contrast, TM treatment increased *atf-4* mRNA levels by only 1.5-fold (Fig. 1d, Extended Data Fig. 1g). Treatment with alpha-amanitin, which blocks transcription, prevented TM from increasing the levels of the *atf-4* mRNA but not P*atf-4*(uORF)::GFP fluorescence (Fig. 1c, 1d). Together, the data indicate that the P*atf-4*(uORF)::GFP reporter is regulated post-transcriptionally during ER stress. The endogenous *atf-4* mRNA was expressed at steady levels during development and ageing (Extended Data Fig. 1h). By contrast, ribosome occupancy was decreased on its uORFs compared to the coding region as larval development progressed, with the lowest levels apparent during the L4 stage after the body plan has been formed and growth has slowed (Fig. 1e, 1f). This suggests that the endogenous *atf-4* gene is regulated translationally through its uORF region under unstressed conditions, possibly in response to growth-related cues. We conclude that, like mammalian ATF4, *C. elegans atf-4* is regulated translationally and preferentially translated upon conditions of reduced protein synthesis (Extended Data Fig. 1i).

Given that ATF-4 is upregulated by translation suppression during the ISR, we hypothesized that it might be important for lifespan extension by reduced protein synthesis. In *C. elegans*, RNAi-mediated knockdown of various translation initiation factors increases lifespan^7–12^. Notably, the lifespan increases that occurred in response to knockdown of *ifg-1*/eIF4G, *ife-2*/eIF4E, or *eif-1A*/eIF1AY, was abrogated in *atf-4(tm4397)* loss-of-function mutants (Fig. 2a, Supplementary Table 1). Similarly, a low dose of cycloheximide extended the lifespan of wild type (WT) but not *atf-4(tm4397)* animals (Fig. 2b, Supplementary Table 1). Thus, *atf-4* is required for longevity arising from a global reduction in protein synthesis.

**Figure 2.**
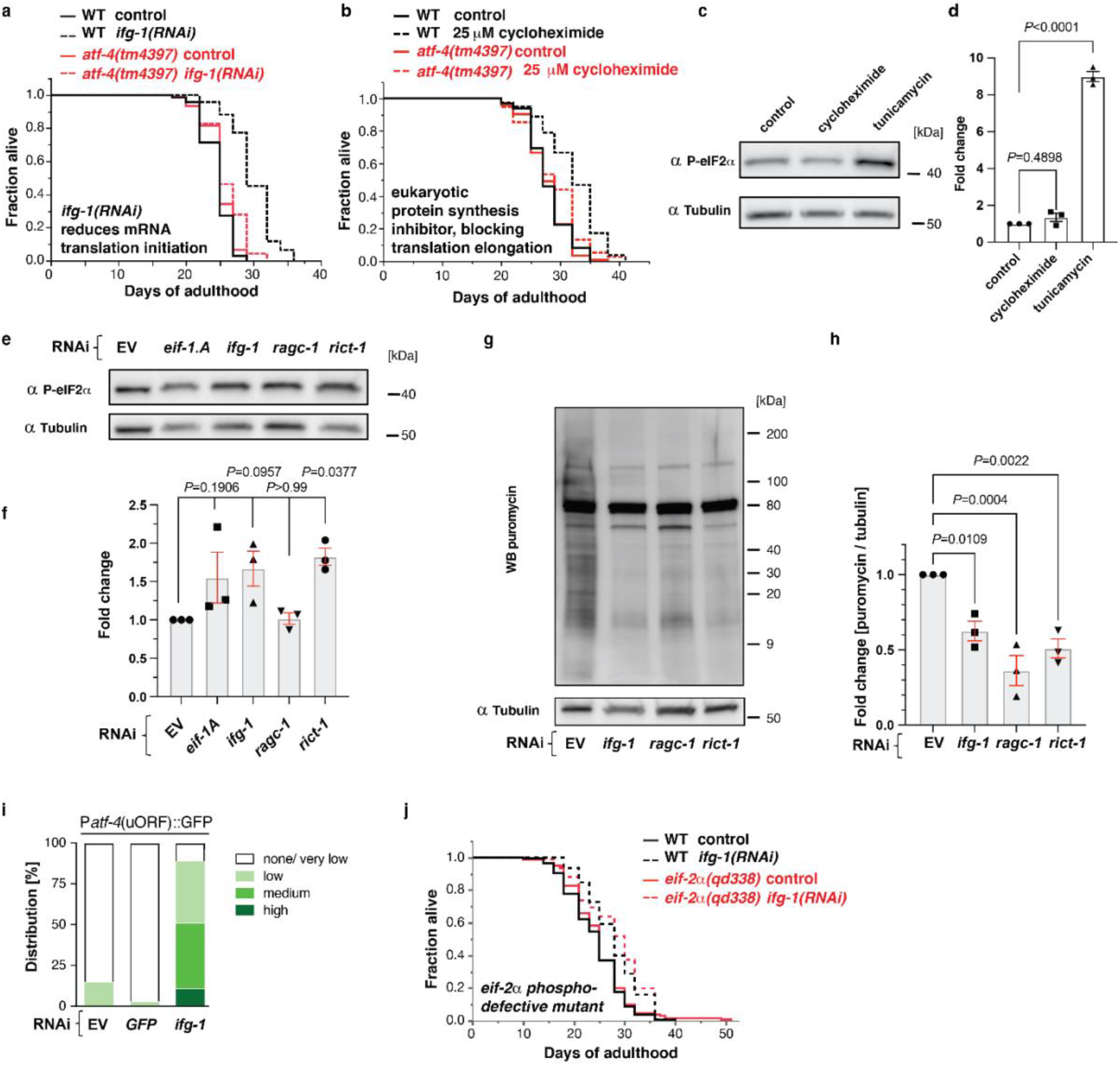
ATF-4 mediates lifespan extension from translation inhibition. **a** Adult-specific knockdown of *ifg-1* extended the lifespan of *WT* animals but not *atf-4(tm4397)* mutants. **b** Adult-specific treatment with 25 μM cycloheximide increased lifespan dependent upon *atf-4*. **c** Representative western blots and quantification (**d**) showing that treatment with 35 μg/ml tunicamycin for 4 hours dramatically increased eIF2α phosphorylation levels in L4 stage animals, while treatment with 7.2 mM cycloheximide for 1 hour did not. One-way ANOVA with post hoc Tukey. **e** Representative western blots and quantification (**f**) showing the effects of adult-specific knockdown of *eif-1.A*, *ifg-1*, *ragc-1*, or *rict-1* on eIF2α phosphorylation levels. One-way ANOVA with Dunnetts’s post-test compared to EV. **g** Representative western blots of puromycin incorporation assay and quantification (**h**) showing that adult-specific knockdown of *ifg-1*, *ragc-1*, or *rict-1* decreased translation. One-way ANOVA with Bonferroni post-test. **i** Quantification of GFP fluorescence showing that adult-specific *ifg-1* knockdown increases expression of P*atf-4*(uORF)::GFP. **j** Adult-specific knockdown of *ifg-1* comparably extended the lifespan of *WT* animals and *eif-2a(qd338)* phosphorylation-defective mutants. For statistics and additional trials in (**a**), (**b**), and (**j**), see Supplementary Table 1.

**Extended Data Figure 2.**
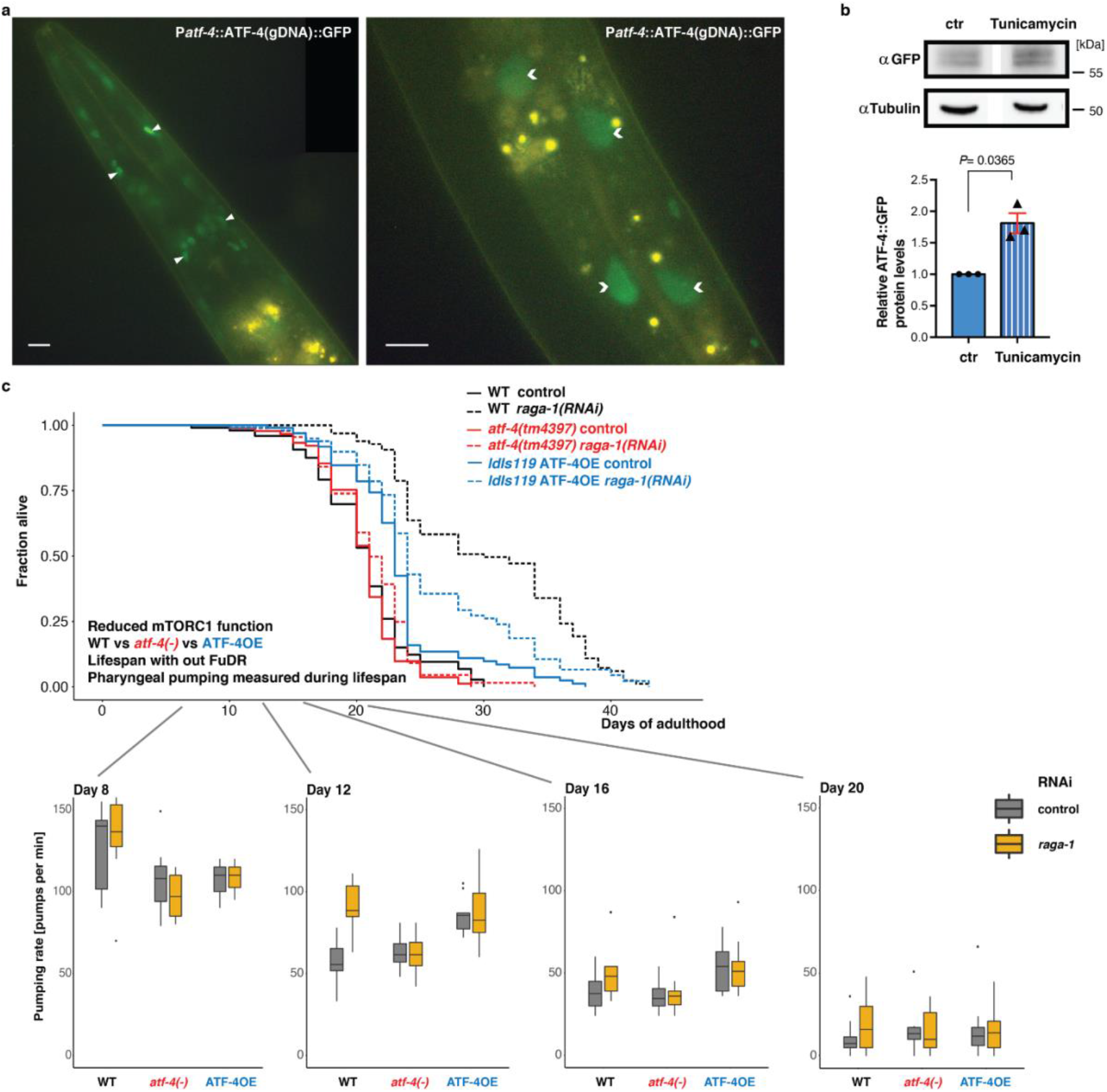
Overexpression of ATF-4 increases healthspan. **a** Representative images showing the expression of ATF-4 in the head (left) and mid-body (right). ATF-4::GFP (*ldIs119*) is displayed in green and found predominantly in nuclei (nuclei of head neurons or glia indicated by arrowheads, intestinal nuclei indicated by chevrons). Yellow puncta are autofluorescent gut granules. 100 x magnification. Scale bar = 10 μm. **b** Western blots and quantification showing ATF-4::GFP levels in day-1-adult transgenic ATF-4OE(*ldIs119*) either treated with DMSO (ctr) or 35 μg/mL tunicamycin for 6 hours. n= 3 replicates. Mean ± SEM. One-sample *t*-test, two-tailed, a hypothetical mean of 1. **c** Pharyngeal pumping measurements across the lifespan comparing WT (N2), *atf-4(tm4397)* mutants, and ATF-4OE(*ldIs119*) treated with either empty vector control RNAi (L4440) or *raga-1(RNAi)*, on plates that do not contain FUdR. See Supplementary Table 2 for raw data.

The increase in ATF-4 translation that occurs during the canonical ISR is induced by translation suppression that is imposed through increased eIF2α phosphorylation^22^, but this might not be the case when translation is reduced by directly inhibiting translation initiation or elongation. Importantly, the low dose of cycloheximide that was sufficient to extend lifespan (Fig. 2b) increased ATF-4 expression (Extended Data Fig. 1f), but not eIF2α phosphorylation (Fig. 2c, 2d). Similarly, depletion of the translation initiation factor *ifg-1*/eIF4G reduced protein synthesis and dramatically increased expression of P*atf-4*(uORF)::GFP, but knocking down either *ifg-1* or *eif-1A* only modestly increased eIF2α phosphorylation (Fig. 2e, 2f, 2g, 2h, 2i). This suggests that treatments that inhibit translation can increase ATF-4 expression without triggering canonical induction of the ISR through eIF2α phosphorylation.

We next investigated whether translation inhibition might increase lifespan independently of this canonical ISR signalling, using a well-characterized eIF2α mutant (*eif2*α*(qd338)*) in which the serine at which inhibitory phosphorylation occurs during the ISR (S49 in *C. elegans*; S51 in mammals) is mutated to phenylalanine, so that eIF2α phosphorylation and ISR induction are blocked^23^. A mutation that prevents phosphorylation of this serine partially suppresses lifespan extension from reduced insulin/IGF-1 signalling, suggesting that ISR signalling is important for longevity in this context^24^. Importantly, the *eif2*α*(qd338)* mutation did not interfere with the *atf-4*-dependent increase in lifespan that was seen with *ifg-1* knockdown (Fig. 2j). We conclude that canonical ISR signalling through eIF2α phosphorylation is not necessarily required for translation inhibition to induce preferential ATF-4 translation or increase lifespan through ATF-4.

### ATF-4 mobilises canonical pro-longevity mechanisms

In *C. elegans*, a limited number of transcription factors have been identified that increase lifespan when overexpressed, including DAF-16/FOXO, HSF-1/HSF1, and SKN-1/NRF^1, 25^. These evolutionarily conserved regulators are generally associated with enhancement of protective mechanisms such as stress resistance, protein folding or turnover, and immunity. To determine whether ATF-4 can actually promote longevity, as opposed to being required generally for health, we investigated whether an increase in ATF-4 levels might extend lifespan. Transgenic ATF-4-overexpressing animals (ATF-4OE) exhibited nuclear accumulation of ATF-4 in neuronal, hypodermal, and other somatic tissues under unstressed conditions (P*atf-4*::ATF-4(gDNA)::GFP; Extended Data Fig. 2a). TM treatment doubled ATF-4 protein levels (Extended Data Fig. 2b, Supplementary Data File 1), indicating that this ATF-4 transgene responds to environmental stimuli. Importantly, ATF-4 overexpression increased lifespan by 7-44% across >10 independent trials, which included two experiments without 5-Fluoro-2’deoxyuridine (FUdR) and analysis of two independent transgenic lines (Fig. 3a, Supplementary Table 1). ATF-4 overexpression also prolonged healthspan (Fig. 3b, Extended Data Fig. 2c, Supplementary Table 2). We conclude that the elevated activity of the ATF-4 transcriptional program is sufficient to extend lifespan and promote health.

**Figure 3.**
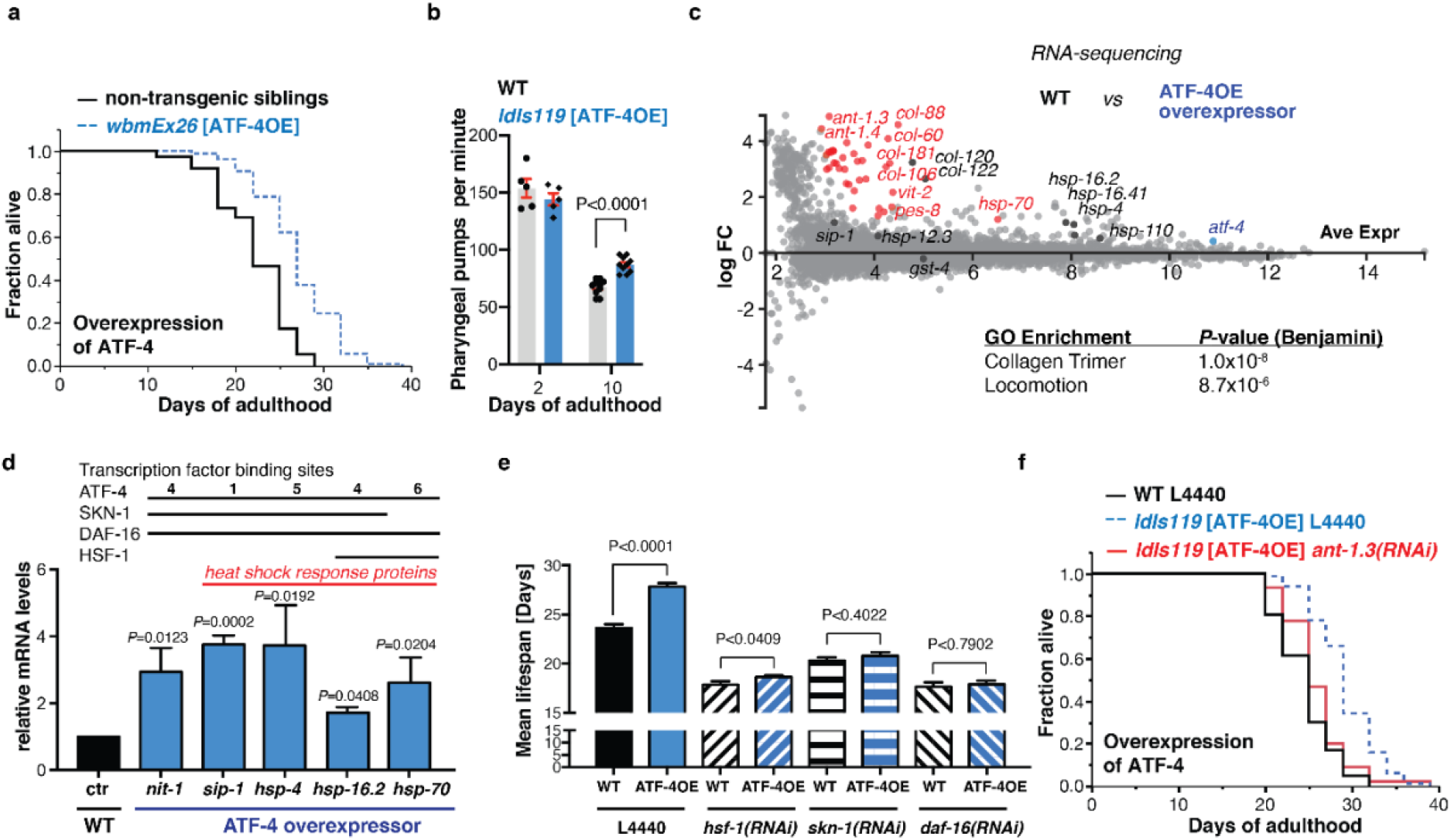
ATF-4 overexpression is sufficient to increase lifespan. **a** Transgenic animals (*wbmEx26* [P*atf-4*::ATF-4(gDNA)::GFP]) that overexpress ATF-4 (ATF-4OE) live longer compared to their non-transgenic siblings. **b** Pharyngeal pumping rate is similar at day 2 of adulthood between ATF-4OE (*ldIs119* [P*atf-4*::ATF-4(gDNA)::GFP]) and WT, but higher in ATF-4OE at day 10 of adulthood, suggesting an improved healthspan. For the complete time-course of pharyngeal pumping rate during ageing, see Supplementary Table 2. Mean ± SEM. Unpaired two-tailed *t*-test. **c** MA (log ratio and mean average)-plot of RNA sequencing analysis comparing ATF-4OE(*ldIs119*) to abs log FC relative to WT. In red, highlighted genes with FDR < 0.1 and log FC > 1 compared to WT. In black, genes with FDR > 0.1. Details are in Supplementary Table 3. **d** Validation by qRT-PCR of genes differentially expressed in ATF-4OE(*ldIs119*), using two new independent biological samples of over 200 animals each. Mean ± SEM. *P* values relative to WT determined by one sample *t*-test, two-tailed, hypothetical mean of 1. The numbers of ATF4 binding sequences (-TGATG-)^27, 28^ are indicated in Supplementary Table 4. The DAF-16 and SKN-1 transcription factor binding sites are based on chromatin immunoprecipitation ChIP data from www.modencode.org (Supplementary Table 5). **e** Longevity conferred by ATF-4OE(*ldIs119*) is abolished by knockdown of *hsf-1, skn-1, or daf-16*. Mean ± SEM. **f** The mitochondrial ATP translocase *ant-1.3* is required for ATF-4 overexpression-induced longevity. For statistical details and additional lifespan trials in (**a**), (**e**), and (**f**), see Supplementary Table 1.

**Extended Data Figure 3.**
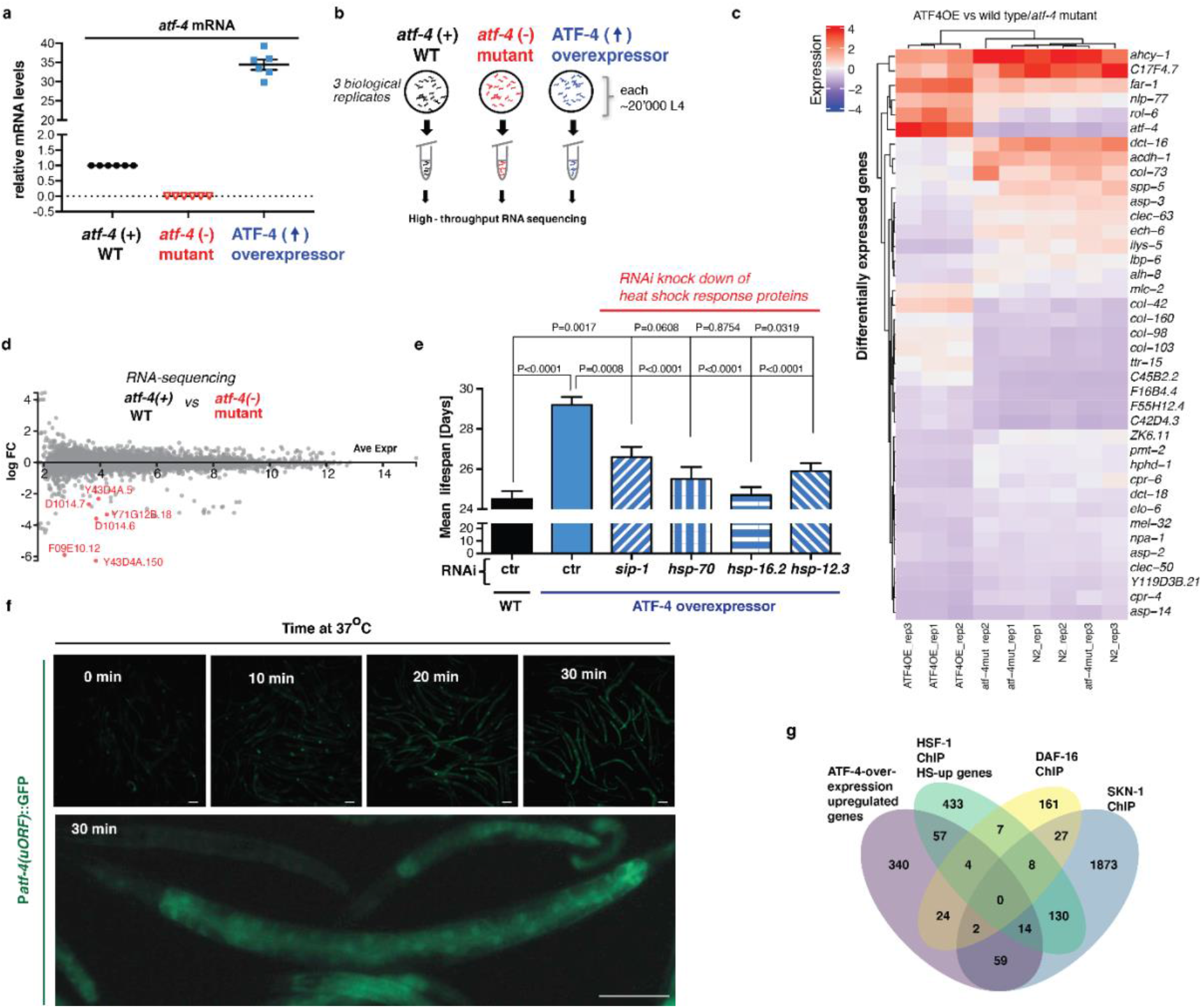
RNA-sequencing reveals transcriptional targets of ATF-4. **a** Quantification of *atf-4* mRNA expression levels in *atf-4(tm4397)* mutants (*atf-4*(-)) and ATF-4OE(*ldIs119*) relative to wild type (*atf-4*(+) WT) animals by qRT-PCR. The same samples were used for RNA sequencing. n=3 independent biological replicates of about 20, 000 L4 *C. elegans*. *P* values for both *atf-4(tm4397)* or ATF-4OE(*ldIs119*) are <0.0001 relative to WT determined by one-sample *t*-test, two-tailed, a hypothetical mean of 1. **b** Schematic representation of sample collection for RNA sequencing. See Materials and methods for details. **c** Hierarchical clustering heatmap of the genes that are most differentially regulated in either direction when comparing ATF-4OE(*ldIs119*) to WT and *atf-4(tm4397)* mutants (*atf-4* (-) mutant). As expected, *atf-4* is in the top gene set. The collagen *rol-6* is the co-injection marker for the transgenic *IdIs119*. Independent biological replicates are indicated as “rep#”. For details and raw data see Supplementary Table 3. **d** MA (log ratio and mean average)-plot of RNA sequencing analysis comparing *atf-4(tm4397)* mutants (*atf-4* (-) mutant) to absolute log fold-change (FC) relative to WT. In red, highlighted genes with a false discovery rate (FDR) < 0.1 and abs log FC > 1 to WT. Details in Supplementary Table 3. **e** Longevity conferred by ATF-4 overexpression (*ldIs119*) is blunted by knockdown of *sip-1, hsp-70, hsp-16.2,* or *hsp-12.3*. Mean ± SEM. P values are relative to WT on empty vector RNAi (L4440). For statistical details see Supplementary Table 1. **f** Representative images that heat increases P*atf-4(uORF)*::GFP transgene expression. Bottom panel shows higher magnification. Anterior to the right, ventral side down. Scale bar = 100 μm. **g** Venn diagram showing the overlap of ATF-4 overexpression-upregulated genes with genes that were bound directly by SKN-1, DAF-16, and HSF-1 in chromatin immunoprecipitation (ChIP) studies. For details and references see Supplementary Table 5.

To identify longevity-promoting mechanisms that are enhanced by ATF-4, we used RNA sequencing (RNA-seq) to compare gene expression profiles in *atf-4* loss-of-function or ATF-4OE animals compared to WT under non-stressed conditions (Extended Data Fig. 3a, 3b, Supplementary Table 3). Only a modest number of genes were detectably up- or down-regulated by *atf-4* loss or overexpression, respectively (Fig. 3c, Extended Data Fig. 3c, 3d). Notably, ATF-4 overexpression upregulated several small heat shock protein (HSP) genes that are also controlled by HSF-1/HSF (heat shock factor) and DAF-16/FOXO (Fig. 3d), and are typically induced by longevity-assurance pathways^1^. Each of the ATF-4-upregulated chaperone genes *sip-1*/CRYAA, *hsp-70*/HSPA1L, *hsp-16.2*/HSPB1, and *hsp-12.3/HSPB2* was required for lifespan extension from ATF-4 overexpression (Extended Data Fig. 3e; Supplementary Table 1). Translation of *atf-4* was increased within minutes by a heat shock (Extended Data Fig. 3f), suggesting that ATF-4 functions in tandem with HSF-1/HSF1. Together, the data suggest that ATF-4 enhances proteostasis mechanisms that have been linked to longevity.

Other findings further linked ATF-4 to longevity-associated mechanisms. ATF-4 overexpression increased expression of the cytoprotective gene *nit-1/*Nitrilase (Fig. 3d), a canonical target of the xenobiotic response regulator SKN-1/NRF^25^, along with expression of collagen genes that are typically upregulated by SKN-1/NRF in response to lifespan extension interventions (Fig. 3c)^26^. The 3kb predicted promoter regions of many ATF-4-upregulated genes included not only the binding consensus for mammalian ATF4 (-TGATG-)^27, 28^, but also sites for DAF-16, HSF-1, and SKN-1 (Fig. 3d, Supplementary Table 4, 5). Furthermore, many genes that were upregulated by ATF-4 overexpression had been detected in chromatin IP (ChIP) analyses of these transcription factors (Extended Data Fig. 3g, Supplementary Table 5). Each of those transcription factors is critical for lifespan extension arising from suppression of translation^10, 11^, and we determined that they are also needed for longevity conferred by ATF-4 overexpression (Fig. 3e, Supplementary Table 1). ATF-4 overexpression also robustly upregulated two adenine nucleotide translocase genes (ANT; *ant-1.3* and *ant-1.4,* Fig. 3c). The ANT complex is important for transport of ATP from the mitochondrial space into the cytoplasm, as well as for mitophagy^29^. Both *ant-1.3* and *ant-1.4* were required for longevity by ATF-4 overexpression (Fig. 3f, Supplementary Table 1). Together, our findings suggest that while the transcriptional impact of ATF-4 may seem limited in breadth, it cooperates with other longevity factors to enhance the activity of multiple mechanisms that protect cellular functions, thereby driving lifespan extension.

### ATF-4 increases lifespan through H_2_S production

To identify ATF-4-regulated genes that are conserved across species and might be particularly likely to have corresponding roles in humans, we queried our ATF-4OE vs WT RNA-seq results, and compared the top 200 significantly upregulated *C. elegans* genes against 152 mammalian genes that are thought to be regulated directly by ATF4^28^. Seven orthologues of these genes were upregulated in ATF-4OE (Fig. 4a, Supplementary Table 4), four of which encoded components of the reverse transsulfuration (hereafter referred to as transsulfuration) pathway (*cth-2*/CTH), or associated mechanisms (*glt-1*/SLC1A2, *C02D5.4*/GSTO1, and *F22F7.7*/CHAC1) (Fig. 4b, Supplementary Table 4). The transsulfuration pathway provides a mechanism for utilising methionine to synthesize cysteine and glutathione when their levels are limiting^30^, but the CTH enzyme (cystathionine gamma-lyase, also known as CGL or CSE) also generates H_2_S as a direct product. Underscoring the potential importance of the H_2_S-generating enzyme CTH-2 for ATF-4 function, the levels of its mRNA and protein were each increased by ATF-4 overexpression (Fig. 4c, 4d, 4e, Supplementary Data File 1).

**Figure 4.**
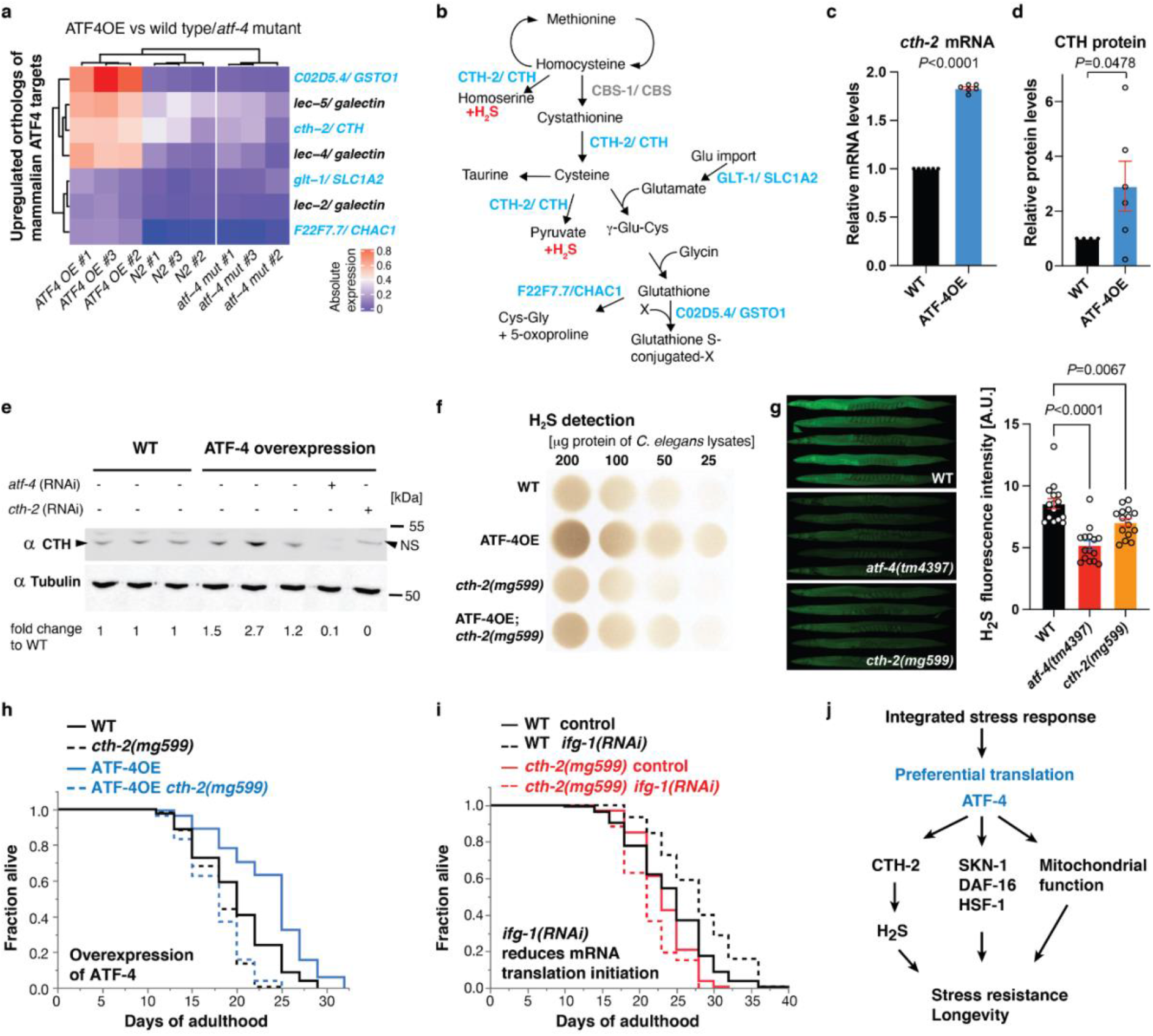
ATF-4 overexpression increases H_2_S levels via cystathionine gamma-lyase, which is required for longevity. **a** Heatmap of gene expression in ATF-4OE (*ldIs119*), wild type (WT), and *atf-4(tm4397)* showing genes whose orthologs are directly regulated by mammalian ATF4 (Details are in Materials and Methods, Supplementary Table 4). Absolute levels of expression were compared. Genes in light blue are predicted to be involved in the transsulfuration pathway, which is shown in (**b**). **c** ATF-4OE(*ldIs119*) showed higher *cth-2* mRNA levels compared to WT by qRT-PCR. n=3 independent biological samples in duplicates (each over 200 L4 worms). Mean ± SEM. *P* values relative to WT determined by one-sample *t*-test, two-tailed, a hypothetical mean of 1. **d** Quantification of CTH protein levels in ATF-4OE(*ldIs119*) compared to WT. n=6 independent biological trials probed in 3 western blots. One-tailed *t*-test. **e** Western blots showing an ATF-4-induced increase in CTH levels was abolished by *atf-4* or *cth-2* knockdown. NS = non-specific band. **f** ATF-4 overexpression increases H_2_S production capacity in a *cth-2-*dependent manner. Additional biological trials are shown in Extended Data Fig. 4d. For H_2_S quantification, see Supplementary Table 12. **g** Representative fluorescent microscopy images and quantification showing that H_2_S levels *in vivo* is decreased in either *atf-4(tm4397)* or *cth-2(mg599)* mutants compared to WT. Data are represented as mean + SEM. *P* values to WT are unpaired *t*-test, two-tailed. **h** Lifespan extension induced by ATF-4 overexpression depends upon *cth-2*. **i** Lifespan extension induced by *ifg-1* knockdown requires *cth-2*. **j** Model for how ATF-4 promotes stress resistance and longevity. For statistical details and additional trials in (**h**) and (**i**), see Supplementary Table 7.

**Extended Data Figure 4.**
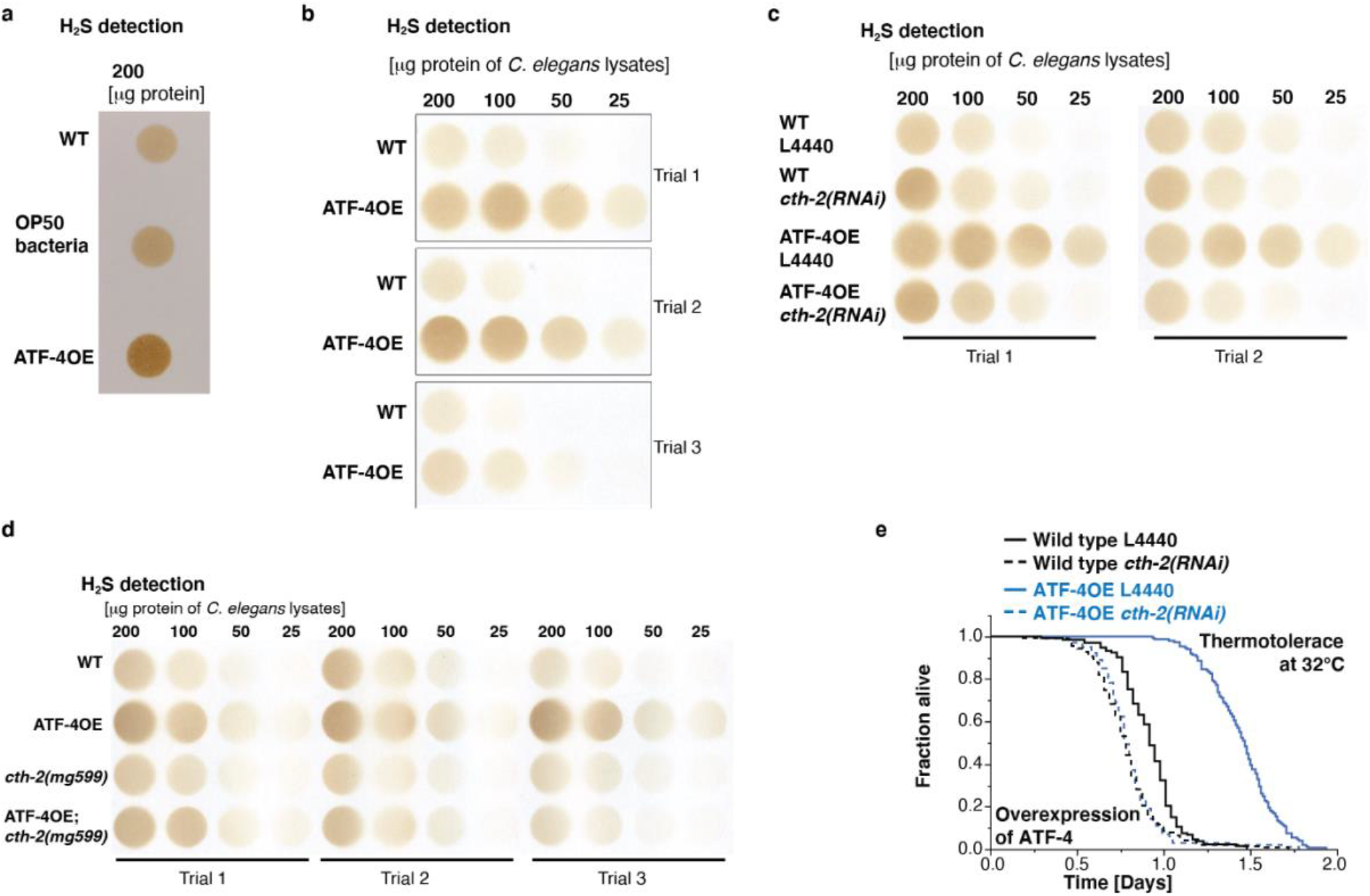
ATF-4 overexpression increases H_2_S production via CTH-2. **a** Measurements of H_2_S production capacity of protein lysates from WT *C. elegans*, the food source OP50 *E. coli* bacteria alone, or ATF-4OE(*ldIs119*) . Because OP50 lysates have the capacity to produce H_2_S, we washed *C. elegans* at least three times or until no bacteria were visible before lysing worms. **b** ATF-4OE(*ldIs119)* exhibited increased H_2_S production capacity compared to WT. n=3 independent biological trials. **c** The increase in H_2_S production capacity in ATF-4OE was reduced by *cth-2* knockdown. **d** Additional trials of Fig. 4f. **e** The increased heat stress resistance deriving from ATF-4 overexpression was suppressed by *cth-2* knockdown. For H_2_S quantification in (**a**)-(**d**), see Supplementary Table 12. For statistical details and additional trials in (**e**) see Supplementary Table 7.

Regulation of amino acid biosynthesis genes is a conserved ATF4 function^13^, and reduced levels of methionine^31^ and higher levels of H_2_S^32, 33^ have each been linked to longevity. We did not detect any differences in the relative abundance of amino acids between ATF-4OE and WT animals (Supplementary Table 6), suggesting that ATF-4 is unlikely to influence longevity by altering amino acid levels. By contrast, ATF-4 overexpression consistently increased H_2_S production capacity in a *cth-2-* dependent manner (Fig. 4f, Extended Data Fig. 4a, 4b, 4c, 4d). Using the fluorescent H_2_S probe MeRho-Az^34^, we also found that H_2_S levels are reduced by mutation of either *atf-4* or *cth-2* (Fig. 4g). Taken together, our data indicate that ATF-4 promotes H_2_S production by acting through CTH-2. The increases in longevity and stress resistance that are conferred by ATF-4 overexpression are abrogated by *cth-2* mutation or knockdown (Fig. 4h, Extended Data Fig. 4e, Supplementary Table 1, 7). Similarly, *ifg-1/*eIF4G knockdown failed to extend lifespan in *cth-2* mutant animals (Fig. 4i). We conclude that the increase in H_2_S production that derived from CTH-2 upregulation is a critical and beneficial aspect of ATF-4 function (Fig. 4j).

An important consequence of increased H_2_S levels is an increase in the proportion of protein cysteine thiols (-SH) that are converted to persulfides (-SSH)^35, 36^. Redox modification at reactive cysteine residues is critical in growth signalling and other mechanisms^36^, and in *C. elegans* thousands of redox-regulated cysteine residues are present in proteins that are involved in translation regulation, lipid and carbohydrate metabolism, stress signalling, and other fundamental biological processes^37^. Under oxidising conditions, these thiols are readily converted to sulfenic acids (-SOH), which is a reversible and in many cases regulatory modification that can proceed to irreversible and potentially damaging oxidized forms (-SO_2_H, -SO_3_H) (Fig. 5a)^36^. H_2_S converts -SOH to -SSH in the process called persulfidation, preventing protein overoxidation and thereby preserving protein functions under stressed conditions (Fig. 5a)^35, 36^. The levels of overall protein persulfidation (PSSH) can be visualised in individual animals by using chemical probes and confocal microscopy^35^. In *C. elegans* PSSH levels are decreased by mutation of the *cth-2* paralog *cth-1*, suggesting that they depend upon a background level of H_2_S produced by the latter^35^. We found that PSSH levels are also reduced by mutation of either *cth-2* or *atf-4*, and are modestly increased by ATF-4 overexpression (Fig. 5b, 5c). Taken together, our results show that ATF-4 acts through multiple mechanisms to promote stress resistance and longevity, and that a CTH-2-driven increase in H_2_S production and persulfidation is an essential aspect of this program.

**Figure 5.**
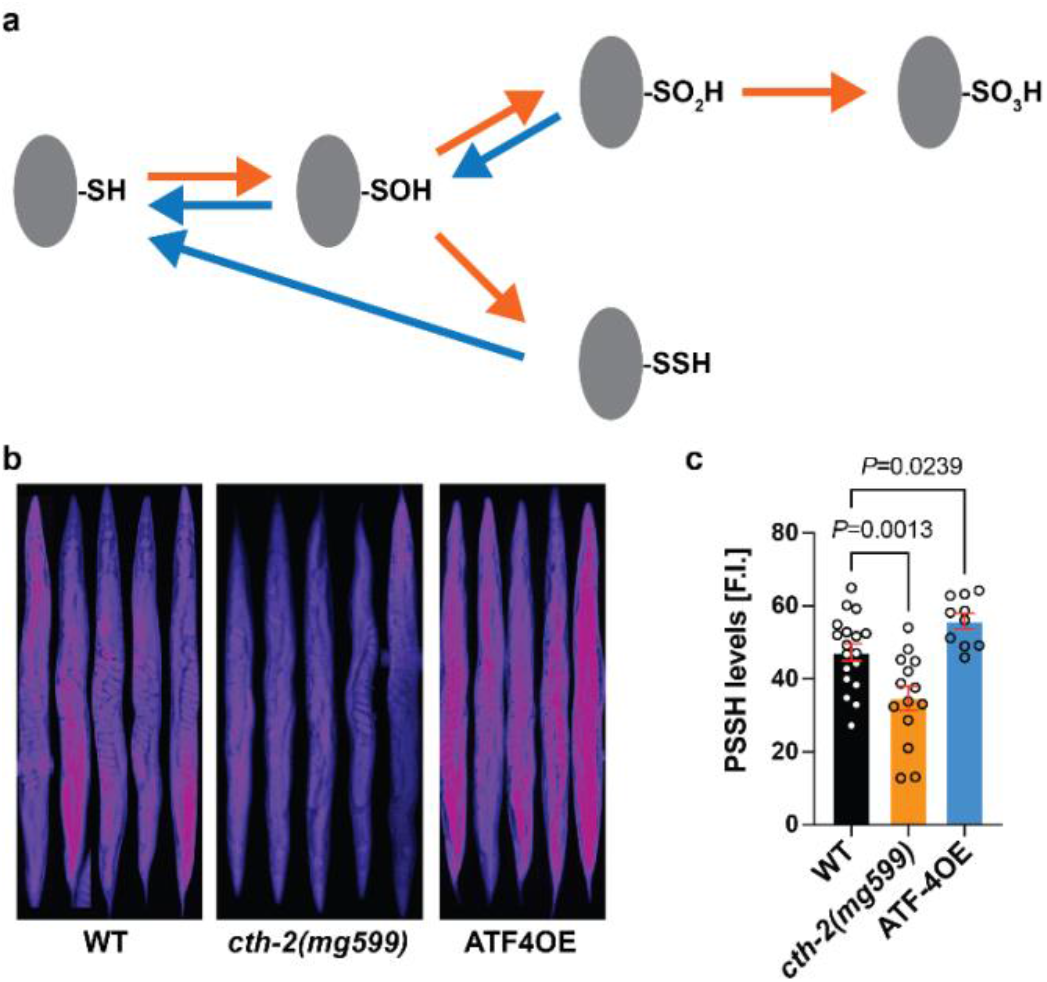
ATF-4 and CTH-2 regulate protein persulfidation levels. **a** Schematic diagram showing that the thiol group (-SH) of reactive cysteine residues in proteins can undertake various redox states. Sulfenylation (-SOH) can be reversed, particularly efficiently through the intermediate of persulfidation (-SSH), but sulfinylation (-SO_2_H) is reversible only within peroxiredoxins and sulfonylation (-SO_3_H) is irreversible^35, 37^. Arrows in orange indicate oxidation processes while those in blue indicate reduction processes. **b** Representative fluorescent images and quantification (**c**) showing that ATF-4OE exhibited higher persulfidation levels, while *cth-2(mg599)* animals exhibited lower global persulfidation levels, compared to WT. Data are represented as mean + SEM. *P* values to WT are unpaired *t*-test, two-tailed.

**Extended Data Figure 5.**
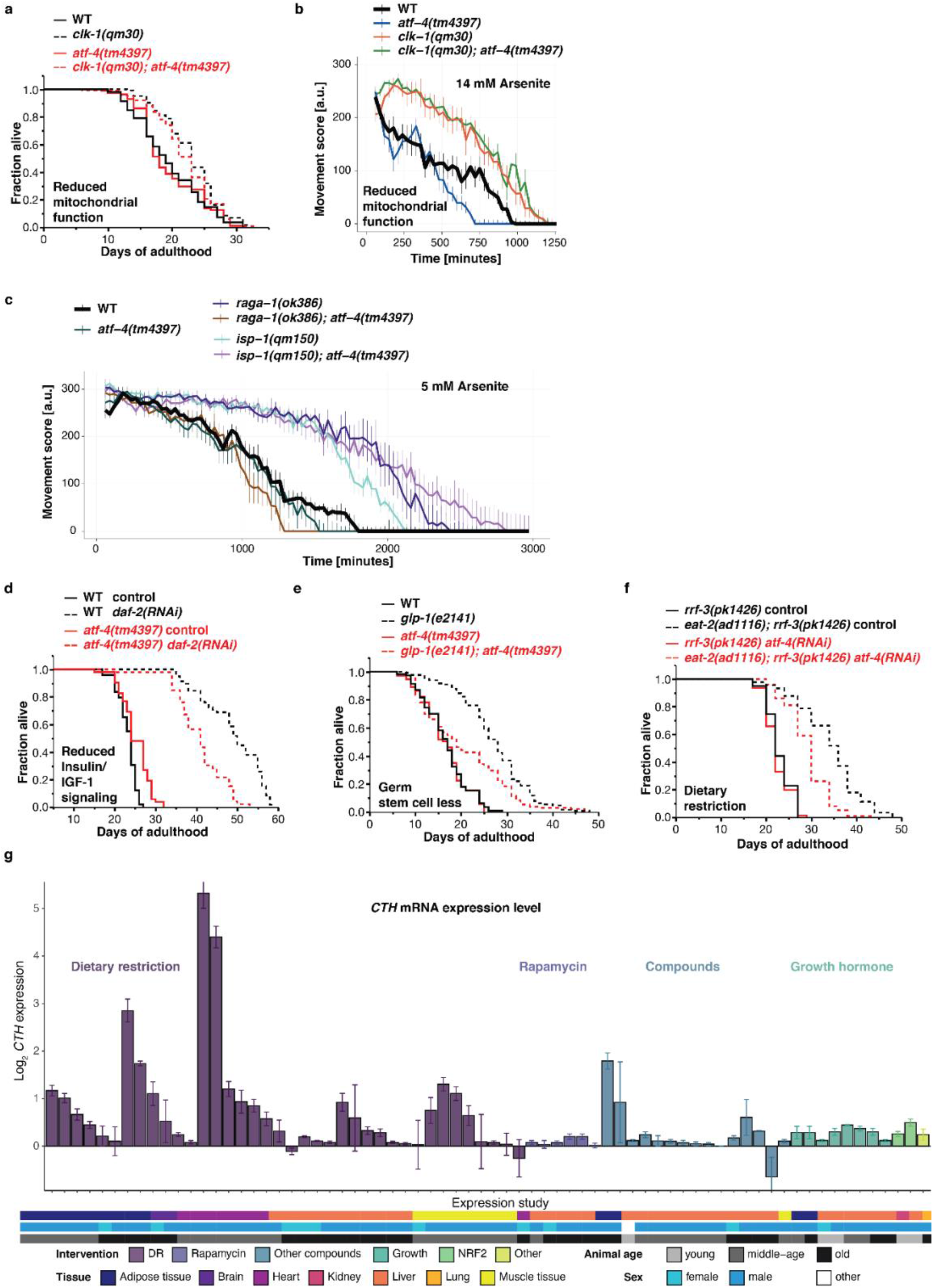
A partial role for ATF-4 in various lifespan extension programs. **a** The longevity of *clk-1(qm30)* animals (impaired mitochondrial function) does not depend on *atf-4*. **b** The increased oxidative stress resistance of *clk-1(qm30)* mutants does not require *atf-4*. **c** Loss of *atf-4* suppresses the oxidative stress resistance of *raga-1(ok386)* mutants (reduced mTORC1 activity), but not that of reduced mitochondrial function mutant *isp-1(qm150)*. **d** Longevity arising from reduced insulin/IGF-1 signalling by adult-specific knockdown of the *daf-2* receptor is partially suppressed by *atf-4(tm4397)* mutation. **e** The *atf-4(tm4397)* mutation partially suppresses the longevity of *glp-1(e2141)* mutants (genetic germline stem cell ablation). **f** Knockdown of *atf-4* partially suppresses the longevity of the genetic DR-related model *eat-2(ad1116)* in the RNAi-sensitized *rrf-3(pk1426)* background. **g** CTH mRNA expression levels in long-lived over control mice, analyzed from publicly available expression datasets (Supplementary Table 11). Data are grouped and colored by interventions and represented as Mean ± SEM. The meta data of the samples are summarized by colored tiles indicating first the tissue of origin, followed by the sex and then the age group of the mice in each experiment. Animals sacrificed before 16 weeks of age were classified as “young”, between 16 to 32 weeks as “middle-aged” and animals above 32 weeks as “old”. If no meta information could be found, it was labelled as “not specified”. For statistical details and additional trials in (**a**), (**d**)-(**f**), see Supplementary Table 1. For statistical details and additional trials in (**b**), (**c**), and see Supplementary Tables 8.

### A partial role for ATF-4 in some lifespan extension programs

Given that *atf-4* is required for lifespan extension in response to reduced translation, we investigated whether *atf-4* and its transcriptional target *cth-2* might be generally required for *C. elegans* longevity. Although ATF4/ATF-4 has been implicated in responses to mitochondrial stress or protein synthesis imbalance^27, 38^, *atf-4* was dispensable for the increases in lifespan or oxidative stress resistance that follow from developmental impairment of mitochondrial function (Extended Data Fig. 5a, 5b, 5c, Supplementary Table 1, 8). The extent of lifespan extension by reduced insulin/IGF-1 signalling or germ cell proliferation was decreased by *atf-4* mutation but did not depend upon *cth-2*, consistent with other transsulfuration components and H2S producers being implicated in the latter pathway (Extended Data Fig. 5d, 5e, Supplementary Table 1)^39^. The ATF-4-CTH-2 pathway of H_2_S production may therefore be fully essential specifically when lifespan extension is driven by suppression of protein synthesis.

We also investigated whether the ATF-4-CTH pathway might be involved in longevity induced by DR, which extends lifespan in essentially all eukaryotes. An increase in H_2_S production capacity has been implicated in mediating some DR benefits in mammals^33^. In *C. elegans*, *atf-4* was not required for lifespan to be extended by a liquid culture food-dilution DR protocol but was partially required for lifespan extension in the genetic DR-related model *eat-2(ad1116)* (Extended Data Fig. 5f, Supplementary Table 1). Consistent with these findings, transsulfuration pathway genes other than *cth-2* are also partially required for *eat-2* lifespan extension^33, 35^, suggesting that in *C. elegans* other pathways than ATF-4-CTH might also increase H_2_S production during DR. In mammals, restriction of sulfur-containing amino acids (methionine and cysteine) acts through ATF4 and CTH to boost endothelial H_2_S levels and angiogenesis^47^, and multiple longevity interventions increase CTH mRNA levels^48^, suggesting a possible role for the ATF4-CTH pathway in DR. Supporting this idea, our bioinformatic analysis revealed that CTH mRNA levels increased in various mouse tissues in response to DR (32/36 profiles), rapamycin (4/6 profiles), or growth hormone insufficiency (8/8 profiles) (Extended Data Fig. 5g, Supplementary Table 11). Therefore, ATF4-induced H_2_S upregulation is likely to be evolutionarily conserved as a contributor to lifespan extension.

### Longevity from mTORC1 suppression is driven by ATF-4 and H_2_S

Because mTORC1 inhibition increases lifespan in part by reducing protein synthesis^2, 5, 6^, we hypothesized that ATF-4 might be activated and required for lifespan extension when mTORC1 is inhibited. The heterodimeric RAG GTPases transduce amino acid signals to activate mTORC1 signalling, and are composed of RAGA-1 and RAGC-1 in *C. elegans*. mTORC1 is required for *C. elegans* larval development^2^, but lifespan can be increased by RNAi-mediated knockdown of either *raga-1* or *ragc-1* during adulthood, or by a partial loss-of-function mutation of *raga-1*^2, 3^. The former strategy allows mTORC1 activity to be reduced without any associated developmental effects. Adulthood RAG gene knockdown reduced protein synthesis (Fig. 2g and 2h), consistent with previous studies of mTORC1^2, 3^, but did not induce eIF2α phosphorylation (Fig. 2e and 2f). Under these conditions P*atf-4(uORF)*::GFP expression was increased robustly even when eIF2α phosphorylation was blocked genetically (Fig. 6a, 6b, Supplementary Table 9). We conclude that ATF-4 is preferentially translated when mTORC1 activity is reduced and translation rates are low, and that this occurs independently of the canonical ISR mechanism of increased eIF2α phosphorylation.

**Figure 6.**
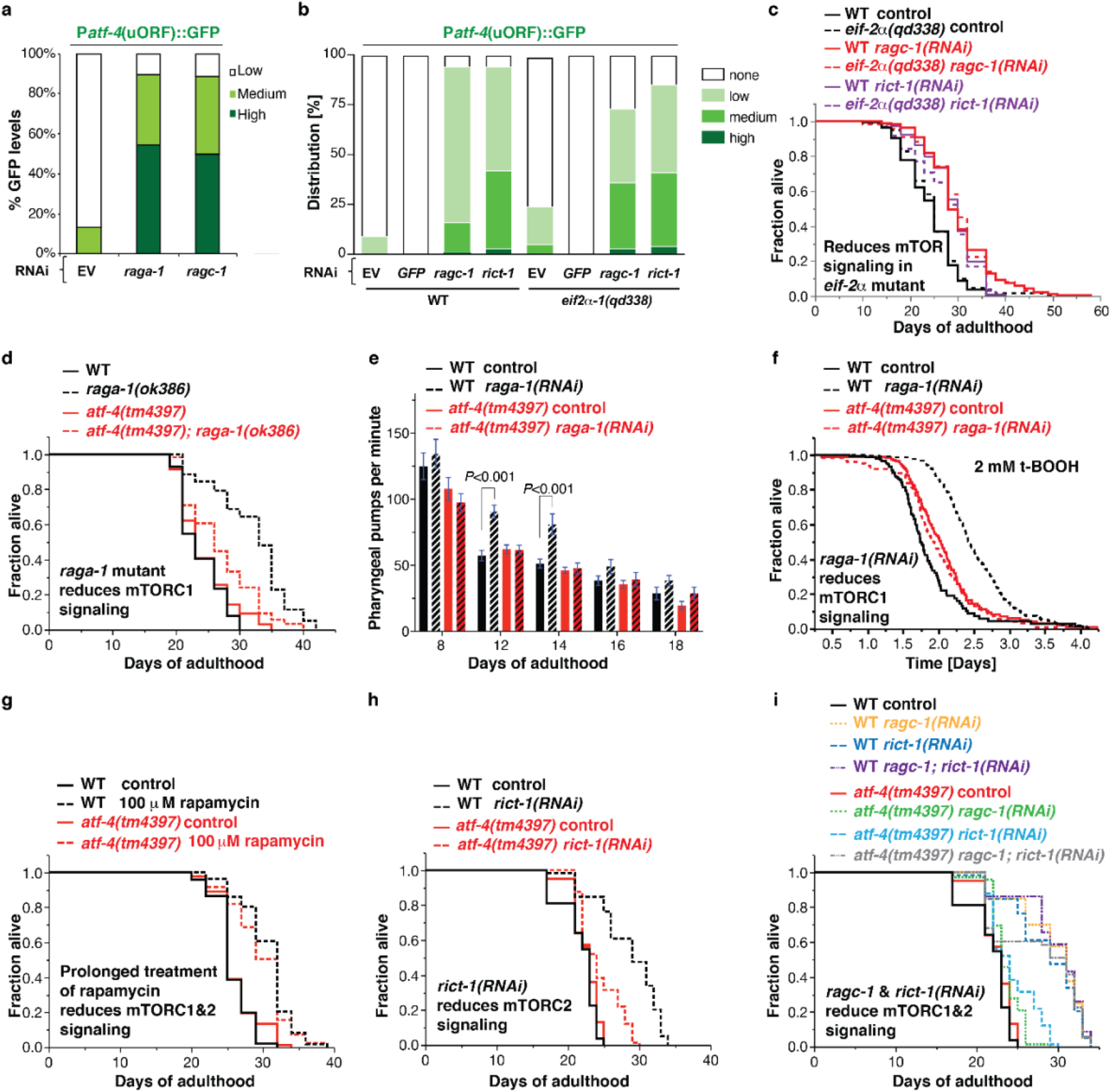
ATF-4 is essential for longevity from reduced mTORC1 activity. **a** Inhibition of mTORC1 by either *raga-1* or *ragc-1* knockdown led to preferential translation of ATF-4. RNAi treatments were initiated at the L4 stage, with GFP intensity scored at day 3 of adulthood. **b** Inhibition of mTORC1 or mTORC2 by knockdown of *ragc-1* or *rict-1*, respectively, leads to preferential translation of ATF-4. Similar effects were observed in WT and *eif2*α*(qd338)* mutants. **c** Post-development knockdown of *raga-1* or *rict-1* extends lifespan in both WT and *eif2*α*(qd338)* mutants. **d** Mutation in *raga-1* increases lifespan in an *atf-4*-dependent manner. **e** Reducing mTORC1 signalling by adulthood specific *raga-1* knockdown improves healthspan dependent upon *atf-4*, as assessed by pharyngeal pumping rate. Mean ± S.E.M. *P* values relative to WT of the corresponding day with One-way ANOVA with post hoc Dunnett’s multiple comparisons test. **f** Adult-specific knockdown *raga-1* increases oxidative stress resistance (2 mM tert-butyl hydrogen peroxide (tBOOH)) in an *atf-4*-dependent manner. RNAi was started at the L4 stage, and stress resistance was measured at day 3 of adulthood with the lifespan machine (See Supplementary Table 10 for details). **g** Rapamycin treatment during adulthood extends lifespan independently of *atf-4*. **h** Adult-specific knockdown of the mTORC2 subunit *rict-1* extends lifespan in an *atf-4*-dependent manner. **i** Adult-specific inactivation of both mTORC1 and mTORC2 increases lifespan independently of *atf-4*. For statistical details and additional trials in (**a**), (**b**), see Supplementary Table 9. For statistical details and additional lifespan trials in (**c**), (**d**), (**g**)-(**i**), see Supplementary Table 1.

**Extended Data Figure 6.**
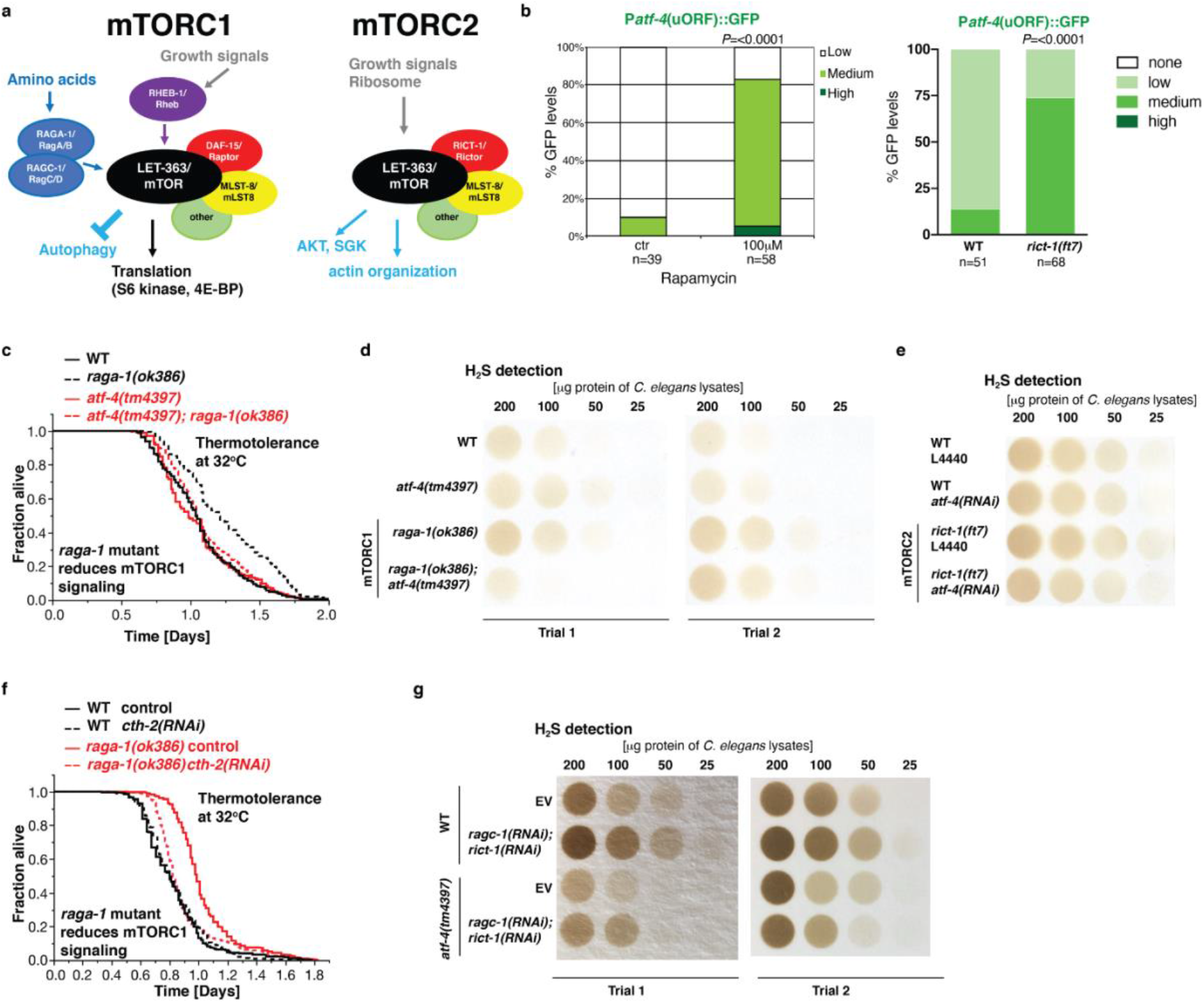
Preferential *atf-4* translation and H_2_S signalling upon reduced mTOR signalling. **a** Schematic diagram showing the composition, regulation, and functions of the two mTOR complexes (mTORC1 and mTORC2) adapted from^6^. **b** Rapamycin treatment (left) or *rict-1(ft7) mutation* (right) leads to preferential translation of ATF-4. Rapamycin treatment was initiated at L4. GFP intensity was scored at day 3 of adulthood. *P* values were determined by Chi^2^ test. **c** Increased heat stress resistance (32°C) of *raga-1(ok386)* mutants depends on *atf-4*. **d** Two additional independent biological trials of Fig. 7a. **e** One additional independent biological trial of Fig. 7b. **f** Increased heat stress resistance (32°C) of *raga-1(ok386)* mutants depends on *cth-2*. **g** H_2_S production capacity assay from *C. elegans* lysates showing that simultaneous knockdown of both *raga-1* and *rict-1* increases H_2_S production capacity in an *atf-4*-independent manner. For statistical details and additional trials in (**c**) and (**f**), see Supplementary Table 7; Quantification of H_2_S assays in (**d**), (**e**), and (**g**) are in Supplementary Table 12.

Importantly, the increases in lifespan extension, stress tolerance, and healthspan that resulted from loss of either *raga-1* or *ragc-1* function required *atf-4* but not phosphorylation of eIF2α (Fig. 6c, 6d, 6e, 6f, Extended Data Fig. 2c, 5c, 6a, Supplementary Table 1-2, 7, 8, 10), consistent with our analyses of P*atf-4(uORF)*::GFP activation. In a single trial, the longevity induced by *raga-1* knockdown was blunted by ATF-4 overexpression, suggesting that too much ATF-4 activity might be harmful (Extended Data Fig. 2c), but this has been observed for some other pro-longevity factors^25^. In summary, the data demonstrate that ATF-4 activation plays an essential role in the benefits of reducing mTORC1 activity

Having determined that *atf-4* is required for mTORC1 suppression to extend lifespan, we were surprised to find that *atf-4* was dispensable for lifespan extension from rapamycin treatment, even though rapamycin increased ATF-4 translational reporter expression (Fig. 6g, Extended Data Fig. 6b, Supplementary Table 1, 9). Notably, mTOR is present in both the mTORC1 and mTORC2 complexes (Extended Data Fig. 6c)^3^. mTORC2 is not as well understood as mTORC1, but it functions in growth signalling and its activation involves binding to the ribosome, suggesting an association with translation regulation (Extended Data Fig. 6c)^40^. Rapamycin mechanistically inhibits mTORC1 activity, but continuous rapamycin treatment also reduces mTORC2 activity by blocking complex assembly^41, 42^, leading us to investigate the possible involvement of *atf-4* in mTORC2 effects. Knockdown of the essential mTORC2 subunit RICT-1 (Rictor) suppressed translation (Fig. 2g, 2h) and increased P*atf-4*(uORF)::GFP expression independently of eIF2α phosphorylation (Fig. 2e, 2f, 6b, Supplementary Table 9). The effects of mTORC2 on *C. elegans* lifespan are complex, but adulthood RNAi knockdown of *rict-1* extends lifespan^2, 6^. Lifespan extension by *rict-1* knockdown required *atf-4* (Fig. 6h, Supplementary Table 1). Interestingly, however, simultaneous inactivation of mTORC1 and mTORC2 by knocking down both *raga-1* and *rict-1* extended lifespan independently of *atf-4* (Fig. 6i, Supplementary Table 1), as occurred with rapamycin treatment, suggesting that when the activity of both mTOR complexes is suppressed the requirement for *atf-4* is relieved.

We investigated whether an ATF-4-mediated increase in H_2_S production is required for lifespan extension arising from inhibition of the mTORC1 or mTORC2 kinase complexes. Genetic inhibition of either mTORC1 or mTORC2 increased H_2_S production capability in an *atf-4*-dependent manner (Fig. 7a, 7b, Extended Data Fig. 6d, 6e). Furthermore, the ATF-4 target gene *cth-2* was required for the increased lifespan of animals with reduced mTORC1 or mTORC2 activity (Fig. 7c, 7d, Supplementary Table 1, 7), suggesting that an ATF-4/CTH-2-mediated increase in H_2_S production is essential for the benefits of either mTORC1 or mTORC2 inhibition. Knockdown of *cth-2* prevented mTORC1 inhibition from increasing stress resistance, further supporting this idea (Extended data Fig. 6f). Interestingly, simultaneous knockdown of *raga-1* and *rict-1* dramatically increased H_2_S production capability, and this was greatly reduced but not eliminated in the *atf-4* loss-of-function mutant (Extended Data Fig. 6g). This result, together with the dispensable role for ATF-4 in longevity under these conditions, suggests that simultaneous inhibition of both mTOR complexes mobilizes mechanisms that increase lifespan independently of *atf-4* and possibly H_2_S.

**Figure 7.**
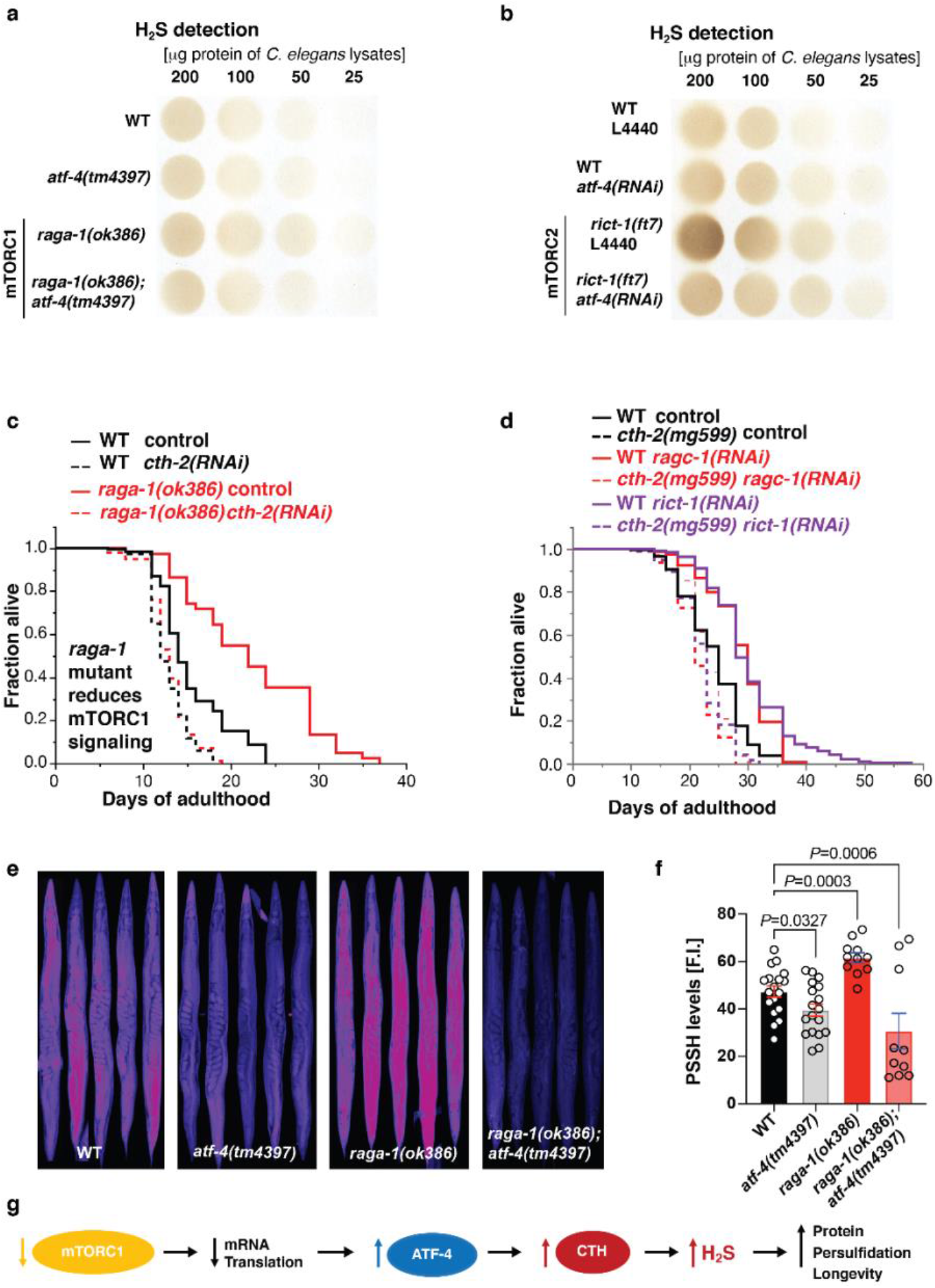
Longevity from mTOR inhibition upregulates H_2_S and requires *cth-2*. **a** Assay of *C. elegans* lysates showing that *raga-1* mutation increased H_2_S production capacity in an *atf-4*-dependent manner. Two additional independent biological trials are in Extended Data Fig. 6d. **b** Assay of *C. elegans* lysates showing that *rict-1* mutation increased H_2_S production capacity in an *atf-4*-dependent manner. An additional independent biological trial is in Extended Data Fig. 6e. **c** Longevity of *raga-1(ok386)* mutants is ablated by *cth-2* knockdown. This particular experiment was performed at 25°C. **d** Longevity induced by adult-specific knockdown of either *raga-1* or *rict-1* depends upon *cth-2*. **e** and **f** Representative images showing persulfidation levels in WT (N2), *cth-2 (mg599)*, *atf-4 (tm4397)*, *raga-1 (ok386)*, and *raga-1;atf-4* mutants. Data are represented as mean + SEM. *P* values to WT are unpaired *t*-test, two-tailed. **g** Inhibition of mTORC1 promotes longevity by increasing ATF-4 expression and stimulating H_2_S production. For H_2_S quantification in (**a**) and (**b**), see Supplementary Table 12. For statistical details and additional lifespan trials in (**c**) and (**d**), see Supplementary Table 1.

The observation that PSSH levels are reduced by lack of either ATF-4 or CTH-2 (Fig. 5b) suggests that the ATF-4/CTH-2 pathway might increase persulfidation in response to interventions that boost H_2_S production through this pathway, including mTORC1 inhibition. Accordingly, in *raga-1* mutant adults PSSH levels were elevated in an *atf-4*-dependent manner (Fig. 7e, 7f). Taken together, our results show that reduced mTORC1 signalling leads to preferential translation of ATF-4, which acts through CTH-2 to promote stress resilience and healthy ageing by increasing H_2_S production, and possibly through the resulting increase in PSSH levels across the proteome (Fig. 7g).

## Discussion

We have identified ATF-4, the transcriptional effector of the ISR, as a pro-longevity factor that can extend *C. elegans* lifespan when overexpressed. Furthermore, conditions that reduce mRNA translation, including mTORC1 inhibition, increase ATF-4 expression without activating canonical ISR signalling that is downstream of eIF2α phosphorylation, and require ATF-4 for lifespan extension. Previous studies revealed that longevity arising from inhibition of translation depends upon preferential translation of protective genes^43^ and increased transcription of stress defence genes^6, 10^. Our new findings link these mechanisms by revealing that preferentially translated ATF-4 cooperates with DAF-16/FOXO, HSF-1/HSF, and SKN-1/NRF to drive protective gene transcription. Viewed alongside the well-documented pro-longevity activity of *S. cerevisiae* Gcn4 (ATF-4 ortholog)^17, 18^ and evidence that ATF-4 levels and activity are increased in long-lived mouse models^44, 45^, our results indicate that ATF-4 has an ancient and broadly conserved function in promoting longevity.

A striking aspect of our findings is that the longevity and health benefits of ATF-4 depend upon activation of its target transsulfuration pathway gene *cth-2*, and the resultant increase in H_2_S production. For many years an understanding of how H_2_S might promote health and longevity proved to be elusive^46^, but recent work has implicated increased PSSH and its salutary effects on the proteome^35^. This modification protects the proteome from the effects of oxidative stress by “rescuing” sulfenylated cysteine residues from the fate of further oxidation (Fig. 5a), declines during ageing, and is increased in other long-lived models^35^. Here we demonstrated that PSSH levels were increased or decreased by ATF-4 overexpression or *cth-2* mutation, respectively, and that mTORC1 inhibition increased PSSH levels through ATF-4 (Figs. 5 and 7). These results provide the first definition of a regulatory pathway through which a pro-longevity intervention increases H_2_S and PSSH levels (Fig. 7g). Although persulfides can be introduced during translation^47^, our results align with previous evidence that H_2_S levels may largely determine the extent of this protective cysteine modification^35^. It was particularly intriguing that the ATF-4/CTH-2 pathway, which boosts H_2_S formation and PSSH levels, was required for lifespan extension from mTORC1 suppression. This ATF-4/H_2_S-induced posttranslational shift in PSSH levels could influence many biological functions, including the activity of redox-regulated signalling pathways^35, 37^, making it of interest to elucidate how these modifications mediate the downstream effects of mTORC1 signalling.

Our data add to the evidence that translation suppression is an essential effector of the longevity effects of mTORC1 inhibition^5, 6^. We were surprised, however, to find that mTORC2 knockdown decreased protein synthesis levels and depended upon ATF-4 for lifespan extension (Figs. 2 and 6). It will be very interesting to elucidate the mechanisms underlying the former observation. It was also surprising that either simultaneous mTORC1 and mTORC2 inhibition or rapamycin extended lifespan independently of ATF-4, suggesting that when both mTOR complexes are inhibited an independent mechanism compensates for lack of ATF-4. An understanding of how this occurs is likely to identify additional mechanisms that promote longevity.

Our evidence that reduced mTORC1 activity promotes longevity by increasing ATF-4 levels contrasts with mammalian evidence that pharmacological mTORC1 inhibition reduces ATF4 translation^20, 48^. However, those findings were obtained in cultured cells in which mTORC1 activity was elevated genetically or by growth factor treatment, a very different scenario from adult *C. elegans in vivo*, in which growth has largely ceased and most tissues are post-mitotic. It seems logical that mTORC1 might increase ATF4 translation under the former conditions, given the importance of mTORC1 for translation of many genes and the need to maintain amino acid levels under conditions of high growth activity. Importantly, it is consistent with our *C. elegans* results that analyses of mouse liver found that ATF4 protein levels and activity are increased in long-lived models that include rapamycin treatment and nutrient restriction^44, 45^, and that mTORC1 hyperactivation (TSC1 deletion) decreases CTH expression and prevents DR from increasing *CTH* mRNA levels^33^. It will be interesting in the future to determine how mammalian mTORC1 influences ATF4 *in vivo* under different conditions, including analysis of tissues with varying rates of growth and levels of mTORC1 activity.

Although inhibition of mTORC1 has received widespread enthusiasm as an anti-ageing strategy, mTORC1 maintains fundamental processes that include protein synthesis, mRNA splicing, and metabolic pathways^2, 3, 48–50^, suggesting that not all effects of mTORC1 suppression are necessarily beneficial. Similarly, although conditions that suppress ISR signalling can promote longevity through effects on selective translation^15^, ISR signalling through eIF2α contributes to lifespan extension from reduced insulin/IGF-1 signalling^24^, and our results demonstrate that the downstream ISR effector ATF-4 is a potent pro-longevity factor. Furthermore, while pharmacological ISR inhibition preserves cognitive functions during ageing by maintaining protein synthesis^14, 16^, our findings suggest that ISR suppression could reduce levels of H_2_S, which has been shown to prevent neurodegeneration^51, 52^. In these and other settings, targeted mobilisation of beneficial mechanisms that are activated by ATF-4, including H_2_S production, might be of promising long-term value. Consistent with this notion, H_2_S confers many cardiovascular benefits in mammals, including a reduction in blood pressure^52, 53^, and patients suffering from vascular diseases show reduced CTH and H_2_S levels^54^, prompting clinical trials of H_2_S-releasing agents for cardiovascular conditions (NCT02899364 and NCT02278276). It could be of considerable value to examine the potential benefits of ATF4 and H2S in various settings, including prevention of ageing-related phenotypes and disease.

### Author contributions

C.Y.E. and T.K.B. conceived the study and designed the experiments. All authors participated in analysing and interpreting the data. C.Y.E., K.P., M.B., S.R.S., C.S., and R.V. performed lifespan assays. C.Y.E., M.B., C.S., and R.V. performed oxidative stress assays. C.Y.E., M.B., and C.S. performed thermotolerance assays. C.Y.E., M.B., K.P., S.R.S., and R.V. scored GFP reporters. C.S., R.E., J.M., A.L., and D.P. performed H_2_S measurement assays. P.L. and C.H. performed Ribo-sequencing analysis. D.P. and M.F. performed persulfidation assays and analysis. C.S. and C.Y.E. analyzed transcription profiles. C.Y.E., C.S., and J.M. performed qRT-PCR. J.M. and R.V. performed the Western bots and puromycin assays. I.M. and W.B.M. generated transgenic strains. C.Y.E. performed all other assays. C.Y.E., T.K.B., and J. M. wrote the manuscript in consultation with the other authors.

### Data availability

The RNA sequencing data in this publication have been deposited in NCBI’s Gene Expression Omnibus and are accessible through GEO Series accession number GSE173799.

## Acknowledgement

We thank Alex Hofer, Anita Goyala, Sara Schütze, Carolin Imse, Julia Rogers, and Lorenza E. Moronetti Mazzeo for help with scoring lifespan, stress, and GFP assays, Michael Steinbaugh for help with the initial analysis of the RNA sequencing data, Stephanie Lin for contributing to earlier stages of this work, S. Mitani and the National BioResource Project for the *atf-4(tm4212* and *tm4397)* alleles, Mike Crowder for the *rars-1(gc47)* allele, Chi Yun and David Ron for the P*atf-4*(uORF)::GFP reporter strain, WormBase for curated gene and phenotype information, Jay Mitchell and Nancy Pohl for comments on the manuscript, and Spalentor and Michael Hall for inspiration. Some strains were provided by the CGC, which is funded by the NIH Office of Research Infrastructure Programs (P40 OD010440). Portions of this research were conducted on the Orchestra High Performance Computer Cluster at Harvard Medical School (NCRR 1S10RR028832-01). Supported by funding from the Swiss National Science Foundation PBSKP3_140135, P300P3_154633, and PP00P3_163898 to C.Y.E. and C.S., and PZ00P3-185927 to A.L., the Leenaards and Novartis Foundation to A.L., ETH Research Grant (ETH-30-16-2) to R.V., the Iacocca Family Foundation to J.M., and NIH R35GM122610 and AG054215 to T.K.B. Part of this research was conducted while Collin Y. Ewald was an Ellison Medical Foundation/AFAR Postdoctoral Fellow.

## Materials and Methods

### Strains

*Caenorhabditis elegans* strains were maintained on NGM plates and OP50 *Escherichia coli* bacteria. LD1499 [P*atf-4*(uORF)::GFP::*unc-54*(3’UTR)] was made by Chi Yun (1.8kb promoter 5’ of *atf-4* including both uORFs into pPD95.75, personal communication with Chi Yun and David Ron)^57^. All strains used are listed in Supplementary Table 13.

### Generation of transgenic lines

Construction of a translational fusion of ATF-4 with GFP. The plasmid pWM48 (P*atf-4*::ATF-4(gDNA)::GFP::*unc-54*(3’UTR)) was generated by introducing the 1.8kb promoter region 5’ of *atf-4* and the *atf-4* genomic sequence into pAD1. This construct was used to generate two independent transgenic lines: *wbmEx26* [pWM48 (P*atf-4*::ATF-4(gDNA)::GFP::*unc-54*(3’UTR), pRF4 (*rol-6(su1006)*)] and *wbmEx27* [pWM48 (P*atf-4*::ATF-4(gDNA)::GFP::*unc-54*(3’UTR), pRF4 (*rol-6(su1006)*)]. UV irradiation was used for integration resulting in *ldIs119* from *wbmEx26* and *ldIs120-1* from *wbmEx27*, which were outcrossed 8-10x against N2.

### Genomic organisation and alignments

Alignment of *C. elegans* ATF-4 (T04C10.4, WBGene00000221, 208 amino acids, www.wormbase.org) with human ATF4 (350 amino acids, P18848; www.uniprot.org) or ATF5 (282 amino acids, Q9Y2D1; www.uniprot.org) sequences was performed by T-COFFEE (Version_11.00.d625267). The *atf-4* genomic representation was made using Exon-Intron Graphic Maker (http://wormweb.org/exonintron) from Nikhil Bhatla. DNA and mRNA sequences were from www.wormbase.org (WS258). For human *ATF4* GenBank BC008090 mRNA sequence was used. The uORFs were predicted with ApE-A plasmid Editor v2.0.50b3. For amino acid alignments T-COFFEE (Version_11.00.d625267) was used.

### Ribosome profiling analysis

Ribosome profiling sequencing data were downloaded from the NCBI Sequence Read Archive (S.R.A.) (http://www.ncbi.nlm.nih.gov/sra/) under accession number SRA055804. Data were analyzed as described^54^: Data analysis was performed with the help of Unix-based software tools. First, the quality of raw sequencing reads was determined by FastQC (Andrews, S. FastQC (Babraham Bioinformatics, 2010)). Reads were then filtered according to quality via FASTQ for a mean PHRED quality score above 30 (http://usegalaxy.org/u/dan/p/fastq). Filtered reads were mapped to the worm reference genome (Wormbase WS275) using B.W.A. (version 0.7.5), and S.A.M. files were converted into B.A.M. files by SAMtools (version 0.1.19). Read counts were then assigned to each gene or non-coding RNA. Library sizes were normalized using the EdgeR software package and the TMM normalization mode. Ribosome profiling was computed separately for each library as the average normalized read counts across all for normalized libraries. Thus, the mapped reads were determined and normalized based on library size for each transcript in each ribosome profiling library. The ATF-4 coverage data for each larval stage (L1, L2 and L4 ) and for the whole transcript (including 5’UTR, exons and 3’UTR) were calculated and exported by SAMtools. The stage-specific averaged coverage data for each gene were plotted using R (https://www.r-project.org).

### Knockdown by RNA interference

RNAi clones were obtained from the Vidal and Ahringer RNAi libraries. RNAi bacteria cultures were grown overnight in LB with carbenicillin [100 μg/ml] and tetracycline [12.5 μg/ml], diluted to an OD600 of 1, and induced with 1 mM IPTG and spread onto NGM plates containing tetracycline [12.5 μg/ml] and ampicillin [50 μg/ml]. For empty RNAi vector (EV) plasmid pL4440 was used as control.

### Manual lifespan assays

Adult lifespan was determined either with or without FUdR as described in Ewald and colleagues^58^. About 100 L4 *C. elegans* per strain were picked onto NGM plates containing OP50 bacteria. The next day, *C. elegans* (day-1-adults) were transferred onto either NGM plates containing 400 µM FUdR and OP50 bacteria or RNAi bacteria. For cycloheximide-treatment lifespan, day-1-adults were transferred on NGM OP50 plates either containing the solvent 0.25% dimethyl sulfoxide (DMSO) alone as a control or cycloheximide (Sigma #C7698) dissolved in 0.25% DMSO. The rapamycin lifespan and liquid DR lifespan assays were performed as described^6, 58^. Animals were classified as dead if they failed to respond to prodding. Exploded, bagged, burrowed, or animals that left the agar were excluded from the statistics. The estimates of survival functions were calculated using the product-limit (Kaplan-Meier) method. The log-rank (Mantel-Cox) method was used to test the null hypothesis and calculate *P* values (JMP software v.9.0.2.).

### Pharyngeal Pumping

Pharyngeal pumping was assessed as described previously^26^. Pharyngeal pumping was determined by counting grinder movements in 45 second intervals while the animals were feeding on the bacterial lawn.

### Puromycin assay

Puromycin incorporation and detection assays were adapted from previous studies^15, 59^. Approximately 500 L4 animals were resuspended in M9 and transferred to NGM plates containing 50 µM FUdR seeded with RNAi bacteria clones. After 3 days, worms were collected in M9 and then transferred to S-basal medium. Worms were incubated with 4 ml S-Basal that contained OP50 and 0.5 mg/ml puromycin for 1 hr. Afterwards, worms were washed with S-basal for three times. Protein extraction and Western blots for puromycin detection were performed as described below.

### Quantitative Real-Time Polymerase Chain Reaction (qRT-PCR) Assays

RNA was isolated with Trizol (TRI REAGENT Sigma), DNAse-treated, and cleaned over a column (RNA Clean & Concentrator^TM^ ZYMO Research). First-strand cDNA was synthesized in duplicate from each sample (Invitogen SuperScript III). SYBR green was used to perform qRT-PCR (ABI 7900). For each primer set, a standard curve from genomic DNA accompanied the duplicate cDNA samples. mRNA levels relative to WT control were determined by normalizing to the number of *C. elegans* and the geometric mean of three reference genes (*cdc-42, pmp-3,* and Y45F10D.4). At least two independent biological replicates were examined for each sample. For statistical analysis, one-sample *t*-test, two-tailed, a hypothetical mean of 1 was used for comparison using Prism 6.0 software (GraphPad).

### RNA sequencing

Three independent biological replicates were prepared by using sodium hypochlorite to harvest eggs and overnight L1 arrest in M9 buffer with 10 μg/ml cholesterol to synchronize *C. elegans*. For each sample, about 20000 *C. elegans* per strain were allowed to develop to the L4 stage under normal growth conditions on NGM OP50 plates at 20°C (about 1000 *C. elegans* per one 10 cm NGM OP50 plate). WT, *atf-4(tm4397)*, and *ldIs119* were grown at the same time for each biological replicate. *C. elegans* were washed from the culturing NGM plates and washed additional 3 times with M9 buffer to wash away the OP50 bacteria. RNA was isolated with Trizol (TRI REAGENT Sigma), DNAse-treated, and cleaned over a column (RNA Clean & Concentrator^TM^ ZYMO Research). The RNA was sent to Dana-Farber Cancer Institute Center core for sequencing (http://mbcf.dfci.harvard.edu). The RNA Integrity Number (RIN) was then assessed by using the Bioanalyzer 2100 (Agilent Technologies), and only samples with a high RIN score were used to prepare cDNA libraries. All nine samples were multiplexed in a single lane. Single read 50 bp RNA-sequencing with poly(A) enrichment was performed using a HiSeq 2000 (Illumina). We aligned the FASTQ output files to the *C. elegans* WBcel235 reference genome using STAR 2.4.0j software (http://code.google.com/p/rna-star/) with an average >80% coverage mapping the reads to the genome. The differential gene expression analysis was performed using Bioconductor (http://bioconductor.org) as described in^60^. Rsubread 1.16.1 featureCounts was used to quantify the mapped reads in the aligned SAM output files. Transcripts with <1 count per million reads were discarded. Counts were scaled to Reads Per Kilobase of transcript per Million mapped reads (RPKM) and deposited as a final output file in (Supplementary Table 3). To analyze the differential expressed genes, we compared *atf-4(tm4397)*, and *ldIs119* to wild type using Degust (http://degust.erc.monash.edu) with the following settings: RPKM with minimum 5 counts using edgeR with a false discovery rate (FDR) of 0.1 and an absolute log fold change (FC) of 1 relative to WT. Results are displayed in MA-plots. Functional annotation clustering was performed with DAVID using high classification stringencies (https://david.ncifcrf.gov).

### Comparison of RNA sequencing data with mammalian ATF4 target genes

The RNA-sequencing data described in the previous section was subjected to differential expression analysis using the limma package (Smyth, Gordon K. “Limma: linear models for microarray data.” Bioinformatics and computational biology solutions using R and Bioconductor. Springer, New York, NY, 2005. 397-420) available in the programming language R (Team, R. Core. “R: A language and environment for statistical computing.” (2013): 201). The 200 most-upregulated genes that were identified by comparison of ATF4 OE to WT and passed a Benjamini-Hochberg adjusted *P*-value threshold of 0.1 were analyzed further. Mammalian ATF4-specific gene targets were obtained from Quiros et al. 2017^27^ and subjected to Ortholist2 to infer *C. elegans* orthologs based on a comparative genomic meta-analysis^61^. The intersection of the most-upregulated genes in our ATF4OE to WT expression analysis and the orthologs of the mammalian ATF4 targets is depicted as a heatmap showing all biological replicates (#1-3) (http://www.bioconductor.org/packages/devel/bioc/html/ComplexHeatmap.html). The *atf-4* mutant samples are shown separately since the displayed genes were selected based on the comparison between ATF4OE and WT. The absolute expression levels are displayed in a blue (low) to white (medium) to red (high) color gradient, with genes indicated as gene names or sequence names if the former is not available. Hierarchical clustering was applied to both genes (rows) and samples (columns). Additional information: GO term enrichment yielded a significant (*P*=0.047, Benjamini-Hochberg corrected) enrichment of the membrane raft compartment (*lec-2, lec-4, lec-5*) while no significant enrichment for GO biological process, GO molecular function, KEGG- or REACTOME pathways were found.

### Analysis of CTH expression levels in mice

Publicly-available expression datasets were analyzed to quantify the change of CTH expression levels in long-lived compared to normal-lived mice. A selected subset of comparisons displaying CTH upregulation in longevity is depicted in Fig. 6b, while the full table is provided in Supplementary Table 11. Microarray datasets and platform information were obtained from GEO (https://www.ncbi.nlm.nih.gov/geo/) followed by mapping probes to their corresponding genes and sequencing information was obtained from SRA (https://www.ncbi.nlm.nih.gov/sra) and processed using Trim Galore (https://www.bioinformatics.babraham.ac.uk/projects/trim_galore/) and Salmon^62^. Datasets were centered and scaled, and subsequently, the mean fold change, as well as its standard error, were computed for the CTH gene.

### Manual thermotolerance assays

Day-1-adults were placed on NGM OP50 plates (maximum 20 *C. elegans* per plate) and placed in an incubator at 35°C. Survival was scored every hour. Animals were classified as dead if they failed to respond to prodding. Exploded animals or animals that moved up on the side of the plate were censored from the analysis. The estimates of survival functions were calculated using the product-limit (Kaplan-Meier) method. The log-rank (Mantel-Cox) method was used to test the null hypothesis and calculate *P* values (JMP software v.9.0.2.).

### Automated survival assays using the lifespan machine

Automated survival analysis was conducted using the lifespan machine described by Stroustrup and colleagues^63^. Approximately 500 L4 animals were resuspended in M9 and transferred to NGM plates containing 50 µM FUdR seeded with OP50 bacteria, RNAi bacteria supplemented with 100 μg/ml carbenicillin, heat-killed OP50 bacteria, or UV-inactivated *E. coli* strain NEC937 B (OP50 ΔuvrA; KanR) containing 100 μg/ml carbenicillin. For oxidative stress assays, tBOOH was added to 2 mM to the NGM immediately before pouring and seeding with heat-killed OP50 bacteria. Animals were kept at 20°C until measurement. Heat and oxidative stress experiments were performed using regular petri dishes sealed with parafilm, while tight-fitting petri dishes (BD Falcon Petri Dishes, 50x9mm) were used for lifespan experiments. Tight-fitting plates were dried without lids in a laminar flow hood for 40 minutes before starting the experiment. Air-cooled Epson V800 scanners were utilized for all experiments operating at a scanning frequency of one scan per 10 - 30 minutes. Temperature probes (Thermoworks, Utah, U.S.) were used to monitor the temperature on the scanner flatbed and maintain 20°C constantly. Animals which left the imaging area during the experiment were censored.

Population survival was determined using the statistical software R^64^ with the survival and survminer (https://rpkgs.datanovia.com/survminer/) packages. Lifespans were calculated from the L4 stage (= day 0). For stress survival assays the moment of exposure was utilized to define the time point zero of each experiment.

### Manual oxidative stress assay (arsenite and tBOOH)

The manual oxidative stress assays were performed as described in detail in the bio-protocol^65^. L4 worms were manually picked onto fresh OP50 plates. The next day, 10– 12 day-one old *C. elegans* were transferred into 24 well plates containing 1 mL M9 Buffer in quadruplicates for each strain and condition (three wells with sodium arsenite (Sigma-Aldrich) and one well M9 as control). For *tert*-Butyl hydroperoxide (tBOOH) stress assay, about 80 L4 *C. elegans* per condition were picked onto fresh RNAi plates. Three days later, 20 day-three-old *C. elegans* were picked onto NGM plates containing 15.4 mM tBOOH (Sigma-Aldrich). The survival was scored every hour until all animas died. Exploded animals were excluded from the statistics. The log-rank (Mantel-Cox) method was used to test the null hypothesis and calculate *P* values (JMP software v.9.0.2.).

### Oxidative stress assay by quantifying movement

*C. elegans* were collected from NGM plates and washed four times by centrifugation, aspirating the supernatant and resuspending in fresh M9 buffer again. After the final wash, the supernatant was removed, and 10 µl of the *C. elegans* suspension pipetted into each well of a round-bottom 96-well microplate resulting in approximately 40 - 70 animals per well. To prevent desiccation, the wells were filled up immediately with either 30 µl M9, or 30 µl M9 containing 6.7 mM or 18.7 mM sodium arsenite yielding a final arsenite concentration of 0, 5, or 14 mM, respectively. Per *C. elegans* strain and conditions, we loaded two wells with M9 as control and six wells with either 5 or 14 mM arsenite as technical replicates. The plate was closed, sealed with Parafilm and briefly stirred and then loaded into the wMicrotracker device (NemaMetrix). Data acquisition was performed for 50 hours, according to the manufacturer’s instructions. The acquired movement dataset was analyzed using the dplyr (https://dplyr.tidyverse.org/reference/dplyr-package.html) and ggplot2 (https://ggplot2.tidyverse.org) R packages.

### H_2_S production capacity assay

The H_2_S production capacity assay was adapted from Hine and colleagues^33^. *C. elegans* were harvested from NGM plates and washed four times by centrifugation and resuspension with M9 to remove residual bacteria. Approximately 3000 animals were collected as a pellet and mixed with the same volume of 2x passive lysis buffer (Promega, E194A) on ice. Three freeze-thaw cycles were performed by freezing the samples in liquid nitrogen and thawing them again using a heat block set to 37°C. Particles were removed by centrifuging at 12000 g for 10 minutes at 4°C. The pellet was discarded, and the supernatant used further. The protein content of each sample was determined (BCA protein assay, Thermo scientific, 23225) and the sample sequentially diluted with distilled water to the required protein mass range, usually 25 - 200 µg protein. To produce the lead acetate paper, we submerged chromatography paper (Whatman paper 3M (GE Healthcare, 3030-917)) in a 20 mM lead acetate (Lead (II) acetate trihydrate (Sigma, 215902-25G)) solution for one minute and then let it dry overnight. The fuel mix was prepared freshly by mixing Pyridoxal 5′-phosphate hydrate (Sigma, P9255-5G) and L-Cysteine (Sigma, C7352-25G) in Phosphate Buffered Saline on ice at final concentrations of 2.5 mM and 25 mM, respectively. A 96-well plate was placed on ice, 80 µl of each sample were loaded into each well and mixed with 20 µl fuel mix and subsequently covered using the lead acetate paper. The assay plate was then incubated at 37°C for 3 hours under a weight of approximately 1 kg to keep the lead acetate paper firmly in place. For analysis, the exposed lead acetate paper was imaged using a photo scanner. H_2_S levels were quantified as the amount of lead sulfide captured on the paper, measured by the integrated density of each well area (Supplementary Table 12). Quantification of H_2_S production was performed by measuring the integrated density using ImageJ, compared to a well next to it that contained no protein for background.

### Detection of H_2_S levels by confocal microscopy

For the quantification of H_2_S levels, worms were synchronized and grown at 20°C on regular NGM plates seeded with OP50-1 until they reached late L4 stage. At this point, 50 animals per strain were transferred to fresh plates containing fluorescent H_2_S probe to develop until the next day. Plates with H_2_S sensor were made by spreading 100 µl of 40 µM MeRho-Az solution (in DMSO)^34^ on the plate surfaces and left to dry for at least 4 h. On the control plates, the same volume of DMSO was spread as a vehicle control. On the day 1 of adulthood, worms were collected by picking, transferred to a tube containing M9 buffer and centrifuged for 1 minute at 400 x g to remove bacteria. Fixation was done in 2% PFA for 20 minutes at 37°C followed by incubation with 4% PFA for 20 minutes at RT with shaking. PFA was removed and worms were washed 3 times with PBS supplemented with 0.01% Triton and twice with PBS. PBS was removed and mounting media (Ibidi, ref. 50001) was added directly to the tube. Worm suspension was transferred to the glass slide, covered with a cover slip and sealed with the nail polish. Samples were recorded on Leica TCS SP8 DLC Digital Light Sheet and Confocal microscope using 10X air objective and VIS (488 nm) laser. Obtained images were first processed with Worm-align open source pipeline for straightening and then analyzed for fluorescence intensity using Cell Profiler software.

### Persulfidation detection by confocal microscopy

Worms were synchronized by putting 15 gravid adults to lay eggs for 2 h. Once the animals reached day 1 of adulthood, they were washed off the plates with M9 buffer, collected into a 1.5 ml tube, and centrifuged for 1 minute at 400 x g. After 3 washes, M9 was removed, and worms were snap frozen in liquid nitrogen. Samples were defrosted by putting the tubes shortly in the water bath, and 200 µl of 5 mM NBF-Cl in PBS supplemented with 0.01% Triton was immediately added to tubes, followed by incubation at 37°C for 1 h with shaking. Worms were washed for 1 minute with ice-cold methanol while mixing, followed by 3 washes with PBS-Triton to remove excess NBF-Cl. Methanol/acetone fixation was performed on ice by incubating the samples in the ice-cold methanol for 5 minutes and then with the ice-cold acetone for 5 minutes. Acetone was removed, and 3 washes with PBS-Triton were performed. Samples were then again incubated with 5 mM NBF-Cl for 30 minutes at 37°C to ensure complete labelling. After the washes were performed in the same order as previously described, worms were incubated with 150 µl of 25 µM DAz-2:Cy-5 click mix for 1 h at 37°C with shaking. For the negative control, worms were incubated with 25 μM DAz-2:Cy-5 click mix prepared without DAz-2. Samples were then washed 2 times for 5 minutes with ice-cold methanol and 3 times for 5 minutes with PBS-Triton to remove excess of the preclick product. DAPI staining was performed by incubating the samples with 300 nM DAPI solution in PBS-Triton for 5 minutes at RT with agitation. After several washes with PBS-Triton and PBS, worms were mounted on glass slide. Samples were recorded on Leica TCS SP8 DLC Digital Light Sheet and Confocal microscope using 10X air objective and 405 nm laser for DAPI, 488 nm laser for NBF-adducts and 635 nm laser for PSSH. Obtained images were first processed with Worm-align open source pipeline for straightening and then analyzed for fluorescence intensity using Cell Profiler software.

### Scoring of transgenic promoter-driven GFP

For P*atf-4*(uORF)::GFP, L4 stage transgenic animals were exposed to chemicals by top-coating with 500 μl of each reagent (alpha-amanitin (Sigma #A2263), cycloheximide (Sigma #C7698), tunicamycin (Sigma #T7765), sodium arsenite (Honeywell International #35000)) or control (DMSO or M9 buffer) onto 6 cm NGM OP50 plates for 30 min to 4 hours, except that rapamycin (LC laboratories) was added to the NGM agar as described^6^. Then GFP fluorescent levels were either (1) scored or (2) quantified. (1) GFP scoring: Transgenic animals were first inspected with a dissecting scope while on still on the plate. GFP intensity was scored in the following categories: 0= none or very low GFP usually corresponding to untreated control, 1= low, 2= medium, and 3= high GFP fluorescence visible. Animals were washed off chemical treated plates, washed again at least twice, placed on OP50 NGM plates and were picked from there and mounted onto slides and GFP fluorescence was scored using a Zeiss AxioSKOP2 or a Tritech Research BX-51-F microscope with optimized triple-band filter-sets to distinguish autofluorescence from GFP at 40x as described^66^. GFP was scored as the following: None: no GFP (excluding spermatheca), low: either only anterior or only posterior of the animal with weak GFP induction, Medium: both anterior and posterior of the animal with GFP but no GFP in the middle of the animal. High: GFP throughout the animal. *P* values were determined by Chi^2^ test. (2) Quantification of GFP fluorescent levels: Animals were washed off reagent-containing plates, washed an additional two times, then placed into 24-well plates containing 0.06% tetramisole dissolved in M9 buffer to immobilize animals. Fluorescent pictures were taken with the same exposure settings (1s) at 10x magnification using an Olympus Cellsens Standard Camera on an inverted microscope. GFP levels were assessed by drawing a line around the animal, measuring mean grey value and using the same area next to it for background using ImageJ. The arbitrary fluorescent value corresponds to mean grey value of the animals minus the background.

### Western blot

About 5000 *C. elegans* (L4 or day-1-adults indicated in figure legends) were sonicated in lysis buffer (RIPA buffer (ThermoFisher #89900), 20 mM sodium fluoride (Sigma #67414), 2 mM sodium orthovanadate (Sigma #450243), and protease inhibitor (Roche #04693116001)) and kept on ice for 15 min before being centrifuged for 10 min at 15’000 x g^67^. For equal loading, the protein concentration of the supernatant was determined with BioRad DC protein assay kit II (#5000116) and standard curve with Albumin (Pierce #23210). Samples were treated at 95°C for 5 min, centrifuged for 1 min at 10’000 x g and 40 μg protein was loaded onto NuPAGE Bis-Tris 10% Protein Gels (ThermoFisher #NP0301BOX), and proteins were transferred to nitrocellulose membranes (Sigma #GE10600002). Western blot analysis was performed under standard conditions with antibodies against Tubulin (1:500, Sigma #T9026), GFP (1:1’000, Roche #11814460001), Cystathionase/CTH (1:2000, abcam #ab151769), Puromycin (1:10’000, Millipore #MABE343), and Phospho-eIF2α (Ser51) (1:1’000, Cell Signaling #9721). HRP-conjugated goat anti-mouse (1:2’000, Cell Signaling #7076) and goat anti-rabbit (1:2’000, Cell Signaling #7074) secondary antibodies were used to detect the proteins by enhanced chemiluminescence (Bio-Rad #1705061). For loading control (*i.e.,* Tubulin) either corresponding samples were run in parallel, membrane was cut if the size of Tubulin and protein of interest were not overlapping, membrane was incubated with loading control after detection of protein of interest on the same blot, or the blot was stripped (indicated in figure legends). For stripping, membranes were incubated for 5 min in acid buffer (0.2 M Glycin, 0.5 M NaCl, pH set to 2 with HCl) and afterwards for 10 min in basic buffer (0.5 M Tris, pH set to 11 with NaOH) and washed with TBS-T before blocking. Quantification of protein levels was determined by densitometry using ImageJ software and normalized to loading control (*i.e.*, Tubulin). Uncropped blots are provided in the Supplementary Data File 1.

